# Metapopulations with habitat modification

**DOI:** 10.1101/2021.05.27.446046

**Authors:** Zachary R. Miller, Stefano Allesina

## Abstract

Across the tree of life, organisms modify their local environment, rendering it more or less hospitable for other species. Despite the ubiquity of these processes, simple models that can be used to develop intuitions about the consequences of widespread habitat modification are lacking. Here we extend the classic Levins’ metapopulation model to a setting where each of ***n*** species can colonize patches connected by dispersal, and when patches are vacated via local extinction, they retain a “memory” of the previous occupant—modeling habitat modification. While this model can exhibit a wide range of dynamics, we draw several overarching conclusions about the effects of modification and memory. In particular, we find that any number of species may potentially coexist, provided that each is at a disadvantage when colonizing patches vacated by a conspecific. This notion is made precise through a quantitative stability condition, which provides a way to unify and formalize existing conceptual models. We also show that when patch memory facilitates coexistence, it generically induces a positive relationship between diversity and robustness (tolerance of disturbance). Our simple model provides a portable, tractable framework for studying systems where species modify and react to a shared landscape.

---

Many interactions between species are realized indirectly, through effects on a shared environment. For example, consumers compete indirectly by altering resource availability [1, 2]. However, the ways that species affect and are affected by their environment extend far beyond the consumption of resources. Across the tree of life, and over a tremendous range of spatial scales, organisms make complex and sometimes substantial changes to the physical and chemical properties of their local environment [3, 4, 5, 6]. Many species also impact local biotic factors; for example, plant-soil feedbacks are often driven by changes in soil microbiome composition [4, 7, 8, 9].

Numerous studies have recognized and discussed the ways such changes can mediate interactions between species, as well as the obstacles to modeling these complex, indirect interactions [5, 7, 10, 11, 12]. In some instances, the effects of environmental modification by one species on another can be accounted for implicitly in models of direct interactions [2, 13, 14], or within the well-established framework of resource competition [12, 15]. But in many other cases, new modeling approaches are necessary.

Because the range of ecosystems where interactions are driven by environmental modification is wide and varied, many parallel strands of theory have developed for them. Examples include “traditional” ecosystem engineers [16, 17, 18, 19, 20], plant-soil feedbacks [4, 7, 21], and chemically-mediated interactions between microbes [5, 12]. Similar dynamics underlie Janzen-Connell effects, where individuals (e.g., tropical trees) modify their local environment by supporting high densities of natural enemies [8, 22, 23, 24], and immune-mediated pathogen competition, where pathogen strains modify their hosts by inducing specific immunity [25, 26, 27, 28]. These last two examples highlight that environmental modification might be “passive”, in the sense that it is generated by the environment itself.

While each of these systems has attracted careful study, it is difficult to elucidate general principles for the dynamics of environmentally-mediated interactions without a simple, shared theoretical framework. Are there generic conditions for the coexistence of many species in these systems? What are typical relationships between diversity and ecosystem productivity or robustness? We especially lack theoretical expectations for high-diversity communities, as most existing models focus on the dynamics of one or two species [4, 7, 16, 17].

To begin answering these questions, we introduce and analyze a flexible model for species interactions mediated by environmental modification. Two essential features of these interactions— which underlie the difficulty integrating them into standard ecological theory—are that environmental modifications are localized in space and persistent in time [10]. To capture these aspects, we adopt the metapopulation framework, introduced by Levins [29], which provides a minimal model for population dynamics with distinct local and global scales. Metapopulation models underpin a productive and diverse body of theory in ecology [30, 31], including various extensions to study multi-species communities [32, 33]. Here, we adopt the simplest such extension, by assuming zero-sum dynamics and an essentially horizontal community [34, 35]. Our modeling framework accommodates lasting environmental modification by introducing a versatile notion of “patch memory”, in which the state of local sites depends on past occupants.

In line with evidence from a range of systems, we find that patch memory can support the robust coexistence of any number of species, even in an initially homogeneous landscape. We derive quantitative conditions for species’ coexistence and show how they connect to existing conceptual models. Importantly, these conditions apply even as several model assumptions are relaxed. We also investigate an emergent relationship between species diversity and robustness, demonstrating that our modeling framework can provide new insight for a variety of systems characterized by localized environmental feedbacks.

## Results

### Model

We consider a community of *n* species inhabiting a landscape composed of many local patches, all linked by dispersal. At any time, each patch may be occupied by a single species or vacant. A patch becomes occupied by colonization from another patch, and is made vacant by local extinction. In contrast to traditional metapopulation models, vacant patches are not homogeneous. Instead, each vacant patch may be in one of *n* states, corresponding to its last resident, and modeling the effects of modification by that resident. We refer to these modifications collectively as patch memory effects. The ability of species to successfully establish in a patch is sensitive to environmental state, and consequently the state of a vacant patch determines the rate at which it is recolonized by any species in the community.

A simple model for community dynamics incorporating these processes (Fig. 1) is:

**Figure 1:**
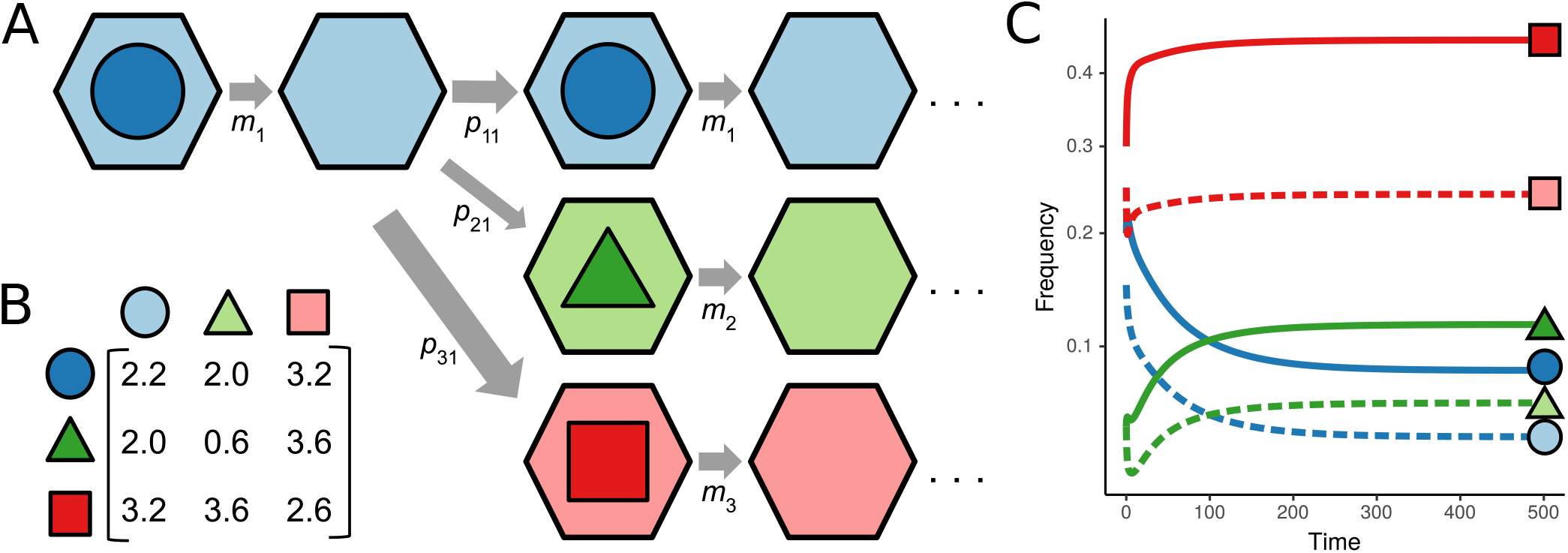
Metapopulation model with patch memory effects. (A) Schematic view of the model. Patches (hexagons) may be occupied by any of *n* (here, 3) species (distinguished by color and shape). Upon local extinction of the resident species, which happens at rate *m*_*i*_, a patch becomes vacant, but retains a “memory” of the last resident (light color). The memory state of a vacant patch determines the rate at which it is (successfully) re-colonized by each species in the community (arrow thickness proportional to rate *p*_*ij*_). Note that colonization events occur at a rate proportional to the frequency of the colonizing species (*p*_*ij*_*x*_*i*_), while local extinction rates are constant per patch. Dynamics for the 3-species community with colonization rates (*P* matrix) given in (B) are shown in (C). The frequency of patches occupied by each species (*x*_*i*_) is shown with solid lines of the corresponding color, while vacant patches in each memory state (*y*_*i*_) are shown with dashed lines. In this example, the colonization rate matrix is symmetric and has exactly one positive eigenvalue, so the community exhibits stable coexistence (see section Symmetric memory effects). In (C), *m* = 1 for all species.

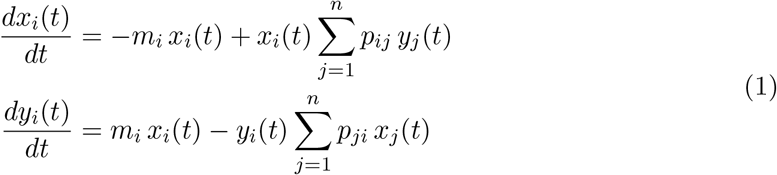

where *x*_*i*_(*t*) is the proportion of patches occupied by species *i* at time *t* and *y*_*i*_(*t*) is the proportion of vacant patches in state *i* (i.e., last occupied by species *i*). The parameters *p*_*ij*_ ≥ 0 specify the rate at which species *i* can colonize patches in state *j*. The local extinction rate for species *i* is given by *m*_*i*_ *>* 0.

This system has close connections to existing ecological models. Eq. (1) reduces to the Levins’ metapopulation model when *n* = 1. Neglecting memory effects, the underlying *n*-species model is conceptually similar to a lottery system [36, 37]. And the full system is closely related to models of infectious disease with multiple strains and partial immunity. As indicated above, when hosts are viewed as patches, with memory effects provided by immunological memory, strain competition is characterized by precisely the kind of dynamics that motivate this study. In fact, our model can be seen as a simplification of the model by Andreasen *et al*. [26], describing a system with short-term immunity in which only the most recent infection is tracked. As widely discussed in the literature on metapopulation dynamics [30] and compartmental models in epidemiology [38], the form of Eq. (1) reflects several standard assumptions: most notably, that the number of patches is large and fixed, and that colonizers or propagules disperse randomly across the landscape. Depending on the context, patches may represent local populations with fast within-patch dynamics, or sites holding a single individual, such as trees in a forest [34].

Patch memory effects in this model are entirely captured by the matrix of species-by-state colonization rates, *P* = (*p*_*ij*_). Rather than track the (potentially many) internal properties of each patch that affect and are affected by resident species, we incorporate them implicitly through these rates. A change in patch state may increase or decrease the rate of colonization by each species, as a net result of factors that facilitate or impede successful establishment. This simplification permits us to model scenarios where the factors mediating patch memory are unknown or complex, and to apply a common modeling framework across systems where these mechanisms differ. Our approach builds on a rich literature modeling metapopulation dynamics in heterogeneous landscapes [39, 40, 41]; however, in our model environmental heterogeneity is generated by the community itself.

We also suppose that the state of a vacant patch depends only the last resident species, and only affects species’ colonization rates, not local extinction rates. Both of these features reflect an underlying assumption that species modify their local environment on a shorter timescale compared to that of extinction. The modifications of each species “overwrite” previous alterations, such that the earlier occupancy history of a patch has negligible effect on current colonization rates. This kind of behavior arises naturally when all species modify a small number of shared environmental features, such as pH or physical structure, and new changes necessarily efface old ones [5, 6]. It is also likely to be a good approximation for many communities where modifications wane without maintenance (see below), resulting in patch memory that is effectively characterized by the most recent occupant [42].

When there is no patch memory (i.e., *p*_*ij*_ = *p*_*ik*_ for all *i, j*, and *k*), the model does not generally admit a biologically feasible equilibrium (an equilibrium with all positive frequencies). The only robust outcome in this case is the eventual extinction of all but one species. However, with patch memory, any number of species may potentially coexist at equilibrium. For a community of *n* species, this full coexistence equilibrium is unique if it exists. Assuming *P* is invertible, the steady-state frequencies for vacant patches are simply ***y***^*∗*^ = *P* ^−1^***m***, where ***m*** = (*m*_1_, *m*_2_ … *m*_*n*_). In general, ***x***^*∗*^, the vector of steady-state frequencies for the *n* species, is obtained by solving an eigenvector problem with no closed form solution (although ***x***^*∗*^ can be written explicitly under special conditions). However, the signs of ***x***^*∗*^, and thus the feasibility of the equilibrium state, can always be determined from ***y***^*∗*^ alone. In SI Model Equations and Coexistence Equilibrium, we show that the conditions 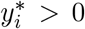 for all *i* and 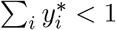 are necessary and sufficient for feasibility.

In this study, we are interested in understanding the long-term dynamics of the model. We first consider cases in which *P* is highly structured, allowing a complete characterization of the long-term dynamics. Then, we go on to examine conditions for coexistence in more general parameterizations of the model.

### Species-specific memory effects

When patch memory effects are species-specific, the colonization rate matrix, *P*, has a very simple structure:

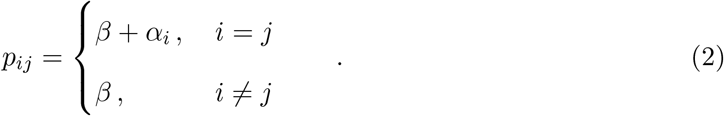

Here, *β* ≥ 0 is the “background” rate of colonization, and *α*_*i*_ expresses the patch memory effects of species *i* on conspecifics. We allow each *α*_*i*_ to be positive or negative, subject to *α*_*i*_ *>* −*β* (ensuring colonization rates are non-negative). This form of *P* models a community where every species modifies and responds to unique properties of the local environment. A number of important and well-studied ecological processes are characterized by this kind of memory effect; for example, local enhancement of specialized natural enemies [22, 23] or specific mycorrhizal symbionts [43] in plant communities, and multi-strain disease dynamics with weak cross immunity [28]. Previous theoretical and empirical studies suggest that memory effects leading to positive intraspecific feedback (*α*_*i*_ *>* 0) should be destabilizing, while negative feedback (*α*_*i*_ *<* 0) provides a source of self-regulation, thereby promoting species diversity [4, 5, 7, 22, 23].

If all species have identical local extinction rates, *m*—as, for example, among demographically equivalent species or when extinction is driven by external disturbance—the steady-state abundance of a species *i* is:

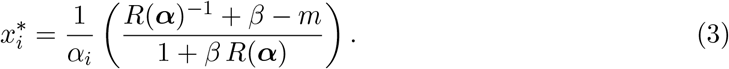

The quantity 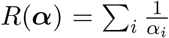 summarizes the net memory effects in the community. Because the factor in parenthesis in Eq. 3 does not depend on *i*, feasibility of the equilibrium (with corresponding ***y***^*∗*^) requires that all *α*_*i*_ have the same sign. In SI Species-Specific Memory Effects, we show that if all *α*_*i*_ *>* 0, the coexistence equilibrium is unstable whenever *n >* 1. When all *α*_*i*_ *<* 0, the equilibrium is instead globally stable, so that coexistence is robust to any perturbation of species frequencies short of extinction. These results align precisely with previous studies that identify stable coexistence with negative environmental feedback.

This highly idealized scenario provides insight into the relationship between diversity and coexistence that extends to systems with more general memory effects. Assuming now that *α*_*i*_ *<* 0 for all *i*, ***x***^*∗*^ will be feasible (and therefore stable) if and only if the common local extinction rate is not too large:

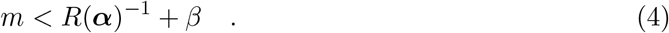

The stability criteria imply that this bound on *m* is strictly increasing as new species are added to the community (Fig. 2A-B). In other words, the model exhibits a positive relationship between diversity and community robustness, defined here as the capacity to tolerate increased disturbance or more marginal environmental conditions (i.e., larger *m*). This effect arises as each new species in the system effectively dilutes the (negative) specific patch memory effects experienced by others. At the same time, however, all species compete for a fixed number of patches. These contrasting effects of diversity are manifest in the relationship between the equilibrium abundance of a focal species, *i*, and *R*(***α***) (i.e., a measure of effective species diversity). When diversity is low, *R*(***α***)^−1^ is very large and negative. Here, 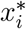 may be increased by the arrival of new species. As more species join the community, however, the beneficial dilution effect of diversity quickly saturates, and 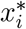 begins to decrease due to the competitive effect of each new species (see, for example, the dynamics of species 1 in Fig. 2A). This transition occurs at the critical value 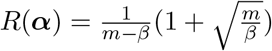.

**Figure 2:**
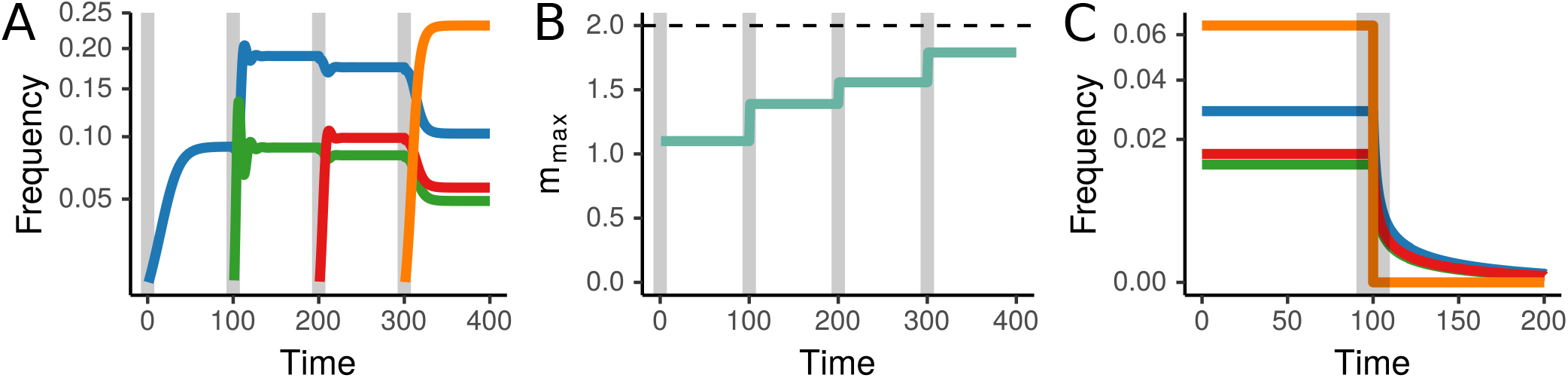
Diversity and coexistence with species-specific memory effects. We consider a pool of 4 species with negative memory effects ***α*** = (−0.9, -1.9, -1.6, -0.4) and *β* = 2. These species coexist stably from any non-zero initial condition, provided that the common extinction rate, *m*, is not too large. The maximum sustainable extinction rate, denoted *m*_max_, is a function of the species present. As successive species join the community (A), *m*_max_ increases (B). Species introductions are marked with vertical gray lines. While *m*_max_ always increases with diversity, eventually saturating at *m*_max_ = *β* (dashed line in (B)), the frequency of individual species may increase or decrease. For example, in (A) the equilibrium frequency of species 1 (blue) increases when species 2 (green) joins the community, but decreases with each subsequent species introduction. (C) The positive relationship between *m*_max_ and species richness means that whole-community collapse is possible when diversity is reduced. Here, all species can coexist with *m* = 1.57, but when species 4 (orange) is removed at time *t* = 100 (vertical gray bar), *m*_max_ drops below *m*, and all species go extinct. In (A), on the other hand, *m* = 1, which permits coexistence at any level of richness. For clarity, unoccupied patch frequencies are not shown.

The results of this section are largely unchanged by variation in local extinction rates among species. When the rates *m*_*i*_ differ, the coexistence equilibrium can become unfeasible; however, the (global) stability of any internal equilibrium point depends only on the signs of *α*_*i*_. Additionally, there remains a well-defined positive diversity-robustness relationship (see SI Species-Specific Memory Effects).

### Symmetric memory effects

Next, we consider the more general class of symmetric memory effects, i.e., *p*_*ij*_ = *p*_*ji*_ for all *i* and *j*. This kind of rate structure may arise whenever patch memory effects depend on some measure of similarity between species. For example, if the local abundances of different predators (pathogens, mutualists, etc.) determine establishment rates, and resident species drive recruitment of their respective predators, then the rates *p*_*ij*_ = *p*_*ji*_ will depend on the number of shared predators of species *i* and *j*. Similarly, for bacterial species with different preferences for environmental pH, and which modify the pH according to their preference, the induced memory effects would be symmetric. This kind of symmetry is considered typical in immune cross-reactivity, as well [44].

For two species, any symmetric matrix can be written in the same form as Eq. 2, so the results of the previous section apply. Assuming feasibility, the two-species equilibrium is globally stable if and only if *p*_12_ = *p*_21_ *>* max (*p*_11_, *p*_22_), regardless of *m*_1_ and *m*_2_. As noted above, this requirement captures the notion that negative feedbacks must predominate in the system for coexistence to be stable. When this condition is met, each species has an advantage colonizing patches last occupied by the other, which generates negative frequency dependence. But how does this intuition generalize to larger communities where patch memory effects cannot be neatly partitioned into intra- and inter-specific components?

In more diverse communities, a different approach is needed to assess stability. Under the assumption of identical local extinction rates (as discussed above) we can study the local stability of the coexistence equilibrium for any *n*. The weaker notion of local stability characterizes the system’s response to small perturbations from equilibrium. For symmetric *P*, we show (SI Symmetric Memory Effects) that the coexistence equilibrium will be locally stable whenever *P* has exactly one positive eigenvalue (Fig. 3). Because *P* is non-negative, it is guaranteed to have a positive eigenvalue, called the Perron eigenvalue. Thus, our stability condition requires that this is the *only* positive eigenvalue of *P*.

**Figure 3:**
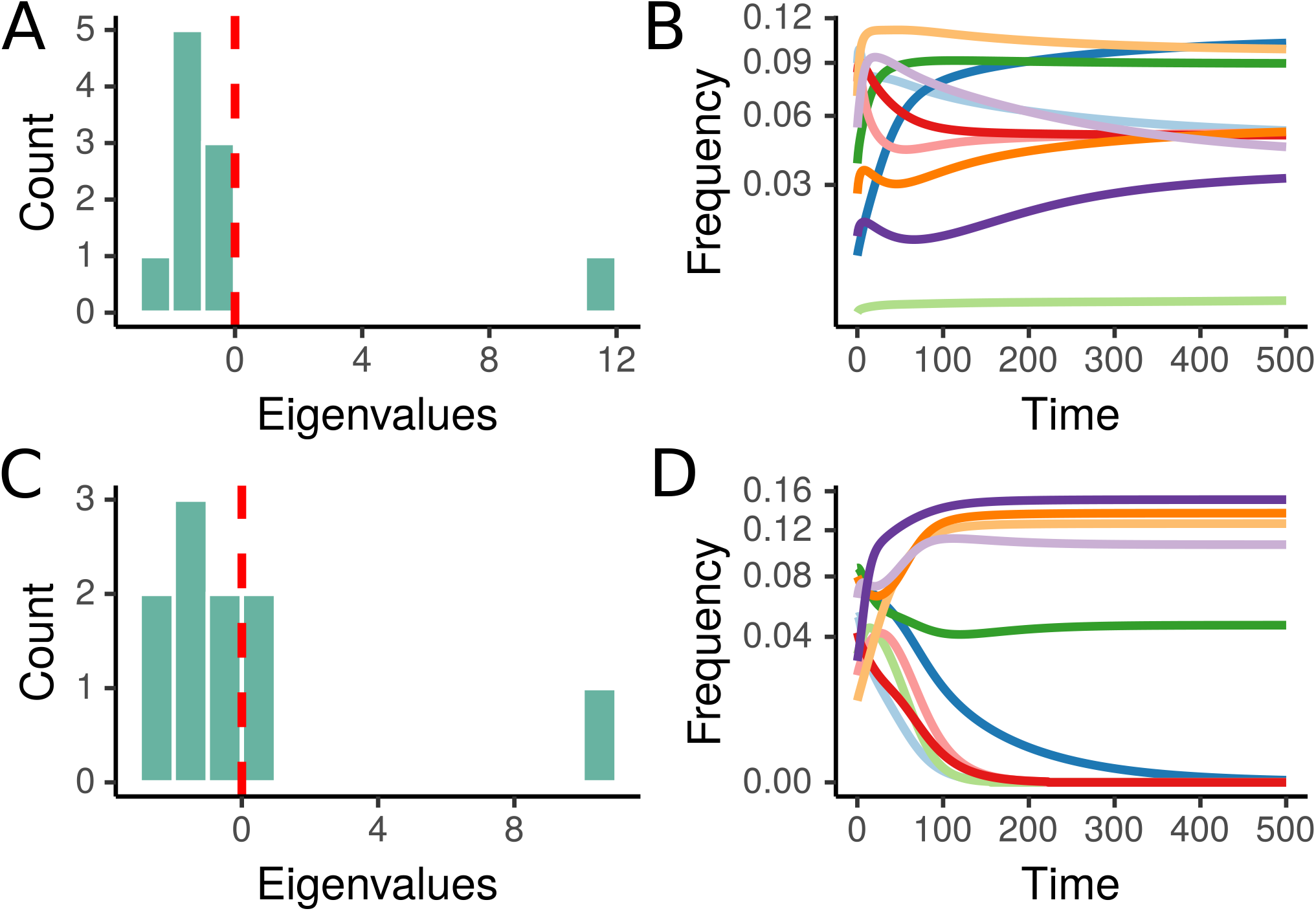
Stability criterion for symmetric memory effects. (A) When the colonization rate matrix *P* has exactly one positive eigenvalue, any feasible equilibrium is locally stable. (B) Coexistence emerges through the dynamics for the 10-species community corresponding to (A). (C) When *P* has any additional positive eigenvalues, the coexistence equilibrium is unstable. (D) For the 10-species community corresponding to (C), 5 species go extinct, despite the existence of a feasible equilibrium. The dynamics prune the community to a subset that meets the coexistence criteria. In (B) and (D), *m* = 1 for all species. For clarity, vacant patch frequencies are not shown. All parameter values can found in the online code [45]

In general, there is no simple characterization of this stability condition in terms of in-equalities between elements of *P*, as was possible for *n* = 2. However, we can find separate necessary and sufficient conditions for a positive symmetric matrix to have exactly one positive eigenvalue [46]. These partial characterizations provide biological intuition for the stability condition. First, a necessary condition: *p*_*ij*_ ≥ min(*p*_*ii*_, *p*_*jj*_) for all *i* and *j*, to have the possibility of coexistence. On the other hand, if 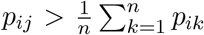 for all *i* ≠ *j*, then a feasible coexistence equilibrium is guaranteed to be stable. Naturally, the second condition is stronger than the first, and in fact implies *p*_*ij*_ *>* max(*p*_*ii*_, *p*_*jj*_). Fig. 1B-C shows an example of a stable system that meets the first (necessary) condition, but not the second. Both conditions place limits on the strength of same-species memory effects (i.e., the magnitudes of *p*_*ii*_) relative to inter-specific effects. In this way, the requirement that *P* has exactly one positive eigenvalue is a natural generalization of the intuitive notion that all species must be at a disadvantage when colonizing patches last occupied by conspecifics.

While this characterization of stable coexistence relies on local stability and identical local extinction rates, extensive numerical evidence suggests that stability is in fact global and unaffected by different *m*_*i*_ (SI Symmetric Memory Effects), as can be proved in the more tractable species-specific case. Expressing equilibrium frequencies (and therefore characterizing feasibility) is also less straightforward for arbitrary *n*. However, the symmetry of *P* means that ***x***^*∗*^ will be proportional to ***y***^*∗*^ (see SI Model Equations and Coexistence Equilibrium). Because ***y***^*∗*^ solves a linear system of equations, a number of techniques may be applied to characterize feasibility and the distribution of species abundances in special cases, such as communities structured by phylogeny or functional groups [47], or when colonization rates are randomly distributed [48]. Additionally, we can ask how diversity affects the “community-wide” feasibility condition, 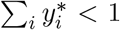. In the previous section, this condition gave rise to a positive relationship between diversity and robustness (i.e., the range of *m* values compatible with coexistence). In SI Symmetric Memory Effects, we prove that the same is true here: For any symmetric system with equal rates *m*, the stability criterion induces a positive diversity-robustness relationship. Specifically, whenever a species is added to a coexisting community and the augmented equilibrium is stable, *m*_max_ must increase. In other words, a coexisting community is always more robust than any of its subsets. While this result alone gives no indication of the strength of the relationship, we show that the robustness benefit of increasing species diversity can be substantial in simulated communities with randomly-distributed memory effects.

### Nonsymmetric memory effects

When patch memory effects are not symmetric, a wide variety of dynamics are possible, including non-equilibrium coexistence. To illustrate the range of outcomes, we consider two particular structures for *P*. We will see that these examples also shed light on typical behaviors for nonsymmetric systems.

Our first example extends the symmetric case to allow variation in dispersal ability. We have so far implicitly assumed that species have equal dispersal rates in defining *p*_*ij*_, which, in practice, represents a composite of two factors: dispersal (ability to reach a new patch) and establishment (ability to successfully establish residence). If each species now has a characteristic dispersal rate, *c*_*i*_, and the establishment rates 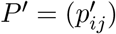 have a symmetric structure, then 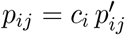 and *P* is *symmetrizable*. While the colonization rate matrix is nonsymmetric, we find that stability is governed solely by the eigenvalues of the establishment rate matrix, *P ′*, according to the stability condition for symmetric matrices (see SI Nonsymmetric Memory Effects).

In our second example, *P* is proportional to a cyclic permutation matrix. Here, patches last occupied by species *i* may only be colonized by species *j*, patches vacated by *j* may only be colonized by *k*, and so on, forming a loop. This intransitive structure is well-known to ecologists, as in the three-species “rock-paper-scissors” dynamics [49]. In our model, this kind of *P* matrix can be seen as a caricature of successional dynamics or facilitation cascades [50], where each species modifies the environment in a way that leads to colonization by the next in the cycle. Strictly interpreted, this structure assumes that the successional process forms a closed loop [21, 51], but these dynamics might also approximate transitive succession in the presence of disturbance [41].

Assuming identical local extinction rates, *m*, and colonization rates, *c*, all species have equal equilibrium frequencies. In contrast to our previous cases, the stability of this equilibrium now depends on the magnitude of *m* (see SI Nonsymmetric Memory Effects). For sufficiently small *m*, the coexistence equilibrium is locally stable and approached by damped oscillations. However, when *n >* 2 there is a threshold value, *m*_*c*_, above which the equilibrium point loses stability and the dynamics approach a stable limit cycle. The amplitude of these cycles increases sharply as *m* increases, making coexistence above *m*_*c*_ tenuous in any real system where environmental and demographic fluctuations would be present. When *m* is sufficiently large, feasibility is lost altogether. For example, the three species community shown in Fig. 4A begins to cycle at *m*_*c*_ = 2*/*9 and loses feasibility at *m*_max_ = 1*/*3. Unlike the symmetric cases, this feasibility threshold decreases as *n* increases, so that diverse communities are actually less robust to elevated local extinction rates.

**Figure 4:**
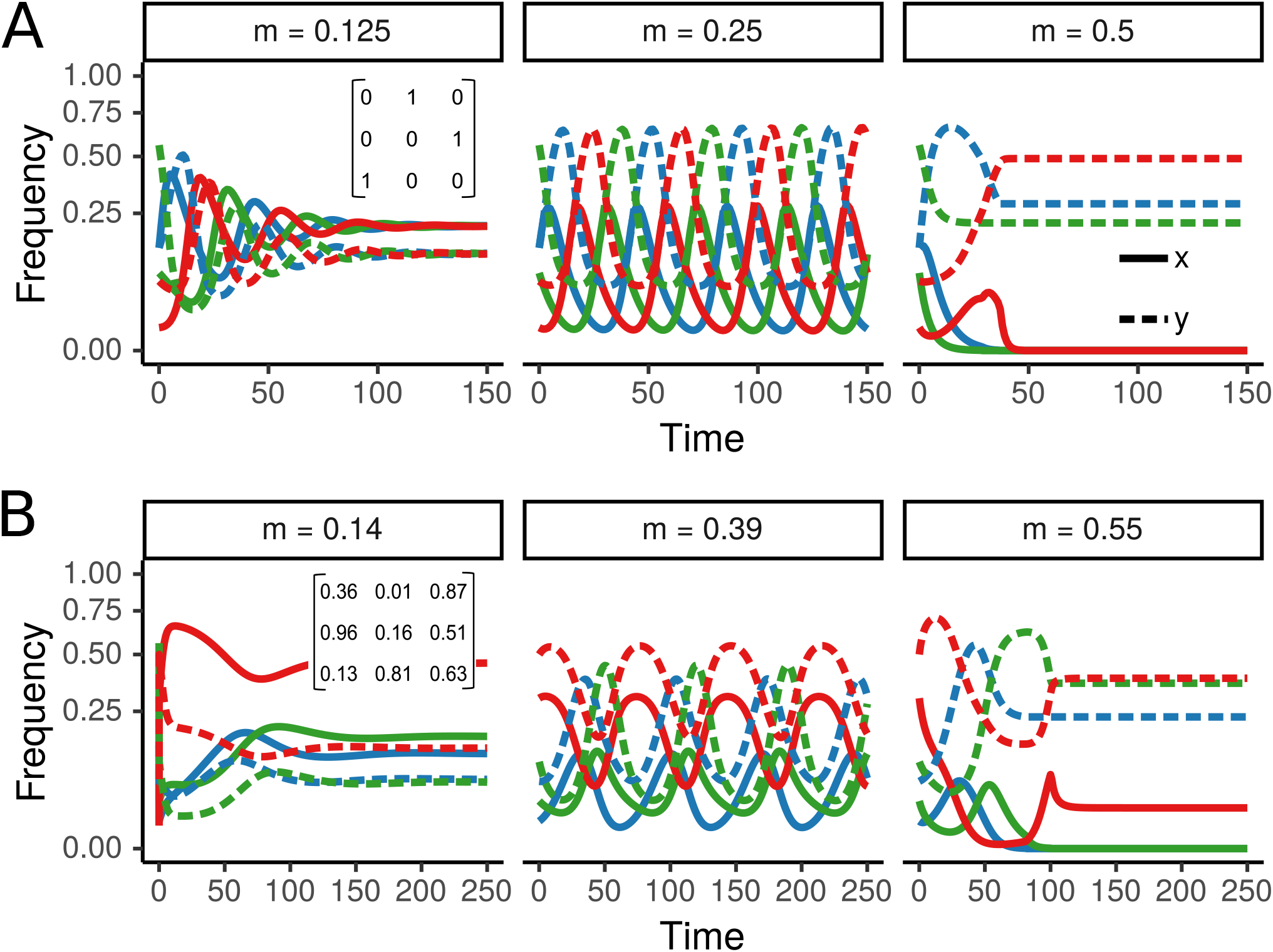
Loss of stability with nonsymmetric memory effects. When *P* is nonsymmetric, the long-term dynamics can depend on the magnitude of local extinction (*m*), even when all species experience equal extinction rates. (A) For the 3-species community with cyclic memory effects, a stable equilibrium exists at low *m*, but gives way to limit cycles at *m* = 2/9. Beyond *m* = 1/3, all species go extinct. (B) This qualitative progression is often observed for nonsymmetric matrices sampled at random, for example the 3-species community shown here. The frequencies of patches occupied by each species (*x*_*i*_) are shown with solid lines, while unoccupied patches in each memory state (*y*_*i*_) are shown with dashed lines.

These qualitative behaviors are shared by many nonsymmetric systems. For randomly-distributed *P*, a progression from stable coexistence, to limit cycles, to species extinctions is often observed as *m* increases and abundances decrease (e.g., Fig. 4B; see SI Nonsymmetric Memory Effects for additional simulations). The frequent appearance of a bifurcation point at high *m* indicates that coexistence mediated by nonsymmetric memory effects is “fragile” in marginal environments. As disturbance increases or environmental quality deteriorates, these systems can collapse sooner than expected on the basis of abundance declines alone (i.e., before feasibility is lost).

### Waning memory

Our model assumes that patch memory persists indefinitely, until a patch is re-colonized. This may be reasonable when the mechanisms mediating patch memory are durable, or when the typical time to re-colonization is short. However, in many systems, we expect patch memory effects to decay in time. For example, nutrient concentrations might re-equilibrate, and physical modifications may erode. This assumption is particularly problematic for modeling pathogen strain competition, where our model is best viewed as an approximation to systems with short-term immunological memory (i.e., immunity well-characterized by the last infection). Clearly this is at odds with the indefinite persistence of memory effects in the absence of new colonization.

To relax the persistent memory assumption, we extended our model to include an additional “naïve” state (see SI Waning Memory Effects). Our extended model mirrors Eq. (1), except that vacant patches of each type *i* decay into a naïve state at a constant rate, *d*_*i*_, simulating waning patch memory. Each species colonizes naïve patches at a constant rate, *c*_*i*_, regardless of the identity of the previous resident.

How do these changes affect the long-term dynamics? For simplicity, we analyze the case where *P* is symmetric, and all species have equal demographic rates *m, c*, and *d*. Under these assumptions, the conditions for coexistence of *n* species are very closely related to those found without waning memory. In particular, the requirement that *P* must have exactly one positive eigenvalue for stability still holds. Now, however, this condition only indicates local, not global, stability of the coexistence equilibrium. This distinction is important because the model can exhibit bistability (discussed below). As before, feasibility of the coexistence equilibrium depends on the signs of ***y***^*∗*^ ∝ *P* ^−1^**1**, in addition to a community-wide condition on the demographic parameters.

The space of outcomes, including global stability of the coexistence equilibrium, bistability, or no feasible equilibrium, is presented graphically in Fig. 5A, and derived in SI Waning Memory Effects. The primary determinant of coexistence is the ratio between *c* and *m*: When *c > m*, the coexistence equilibrium is always feasible, and the only stable fixed point. When *c < m*, coexistence may be impossible (no feasible equilibrium) or contingent on initial conditions (bistability). In this second regime, the coexistence equilibrium is locally stable, but a community-wide Allee effect operates at low species abundances. If the initial fraction of naïve patches is not too large, all species coexist, while if naïve patches are very abundant, all species go extinct.

**Figure 5:**
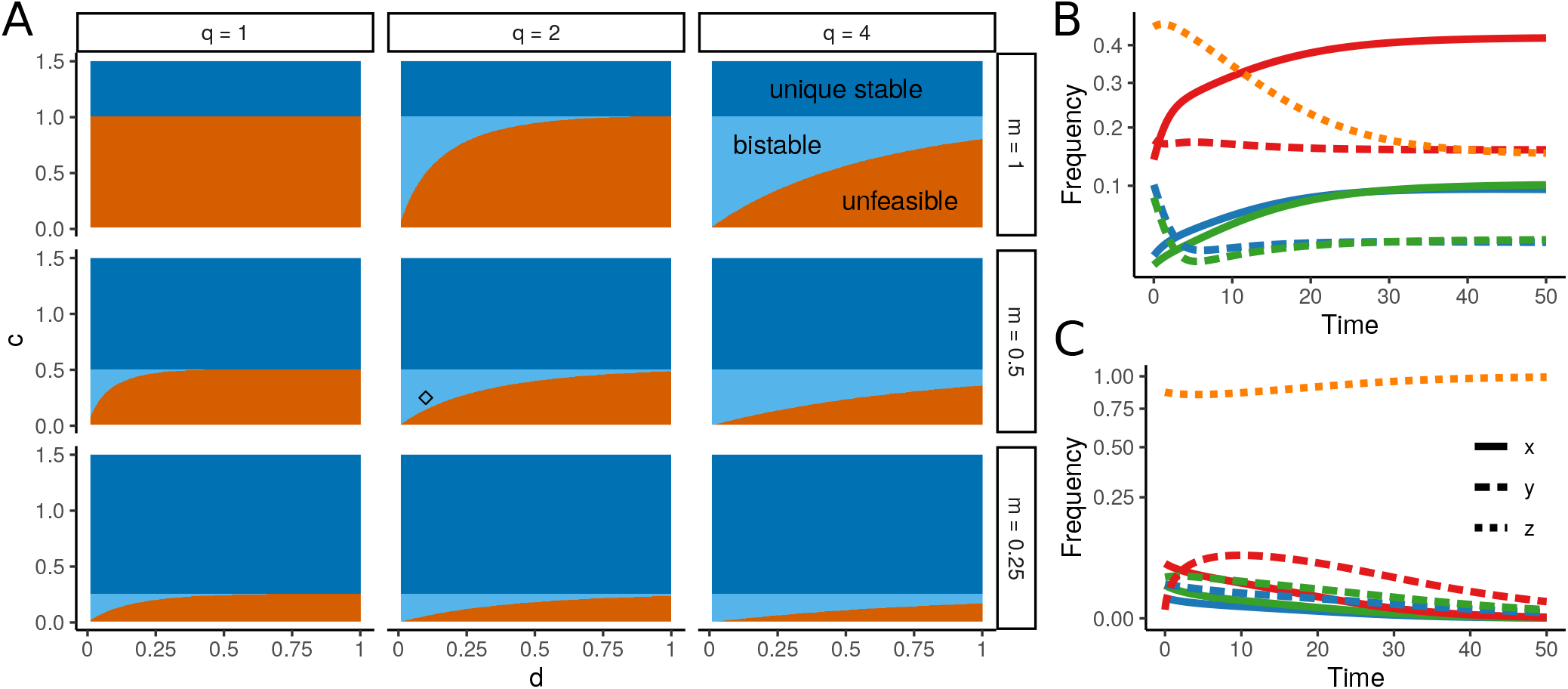
Coexistence conditions for waning (symmetric) memory effects. (A) Assuming *P* has exactly one positive eigenvalue and *P* ^−1^**1** *>* 0 elementwise, the long-term dynamics depend on the relationship between demographic parameters *m* (local extinction rate), *d* (memory decay rate), *c* (colonization rate for naïve patches), and *q* (a statistic quantifying the effective community-wide colonization rate; see main text and SI Waning Memory Effects). When *c > m*, there is a unique, stable coexistence equilibrium for all *d* and *q* (dark blue). When *c < m*, there may be no feasible equilibrium (orange). However, coexistence is possible when *d* is sufficiently small, and *c* and *q* are sufficiently large. In that case (light blue), there is a strong Allee effect. Coexistence is locally stable, but requires that species are not too rare initially. (B) and (C) show two possible outcomes for the parameter combination indicated by the black diamond in (A). (B) When initial frequencies of community members are high enough, all 3 species coexist stably. (C) When naïve patches are initially abundant, the community collapses and all species go extinct. In (B) and (C), the frequencies of patches occupied by each species (*x*_*i*_) are shown with solid lines, unoccupied patches in each memory state (*y*_*i*_) are shown with dashed lines, and naïve patches (*z*) are shown with dotted orange line. The matrix *P* used for this example is from Fig. 1B.

This strong Allee effect is present when *c < m, d* is small, and the net colonization rate of patches with memory (measured by a summary statistic of *P, q* = (**1**^*T*^ *P* ^−1^**1**)^−1^) is sufficiently large. These conditions correspond to an ecosystem where no species is able to colonize naïve patches fast enough to outpace local extinction (*c < m*), but patch memory effects facilitate colonization overall, and memory effects are sufficiently durable to influence the dynamics. Negative feedbacks for each species maintain diversity, while a positive community-wide feedback maintains the growth rates of all species once their frequencies exceed a certain threshold. This picture can become much more complicated if species vary significantly in their rates of memory decay and naïve patch colonization. In this general setting, it is possible for Allee effects to emerge on multiple timescales, leading to more complex multistability.

## Discussion

Species interactions mediated by local environmental feedbacks are ubiquitous in natural systems and arise from many different proximal mechanisms. We presented a highly simplified but flexible model for the dynamics of such communities, with the goal of elucidating general principles that govern coexistence across these disparate settings. Our model recapitulates behaviors observed empirically and in system-specific theory, namely coexistence of many species maintained by negative feedbacks [7, 8, 9, 24, 27]. By virtue of the simplicity of the model, we were able to derive precise conditions for this coexistence, in addition to novel predictions regarding diversity and community robustness, and the emergence of bistability when memory effects are impermanent.

Our central finding, that patch memory effects allow the potential coexistence of any number of species, shares close conceptual similarities with models of facilitation-driven coexistence [52], particularly microbial crossfeeding [53, 54]. In these phenomena and our model, high species diversity can be maintained because community members generate elevated environmental heterogeneity relative to the abiotic background. But in contrast to crossfeeding models, where the number of coexisting species is eventually bounded by the number of distinct resources produced and consumed, our model of patch memory automatically produces the possibility of coexistence no matter how many species are added to the community.

Of course, the possibility of coexistence is no guarantee that it will be realized. We have shown that there are nontrivial—but interpretable—conditions for the feasibility and stability of equilibria in our model. These conditions are unlikely to be met when parameters are drawn at random. However, a number of ecological and evolutionary mechanisms are known that may structure memory effects in way that would promote coexistence [7, 22, 23, 27]. Our model can provide theoretical guidance to help understand these mechanisms and their associated colonization rate structures. For example, there has been significant debate over the degree of enemy specialization needed to produce coexistence through Janzen-Connell effects, which can be modeled (in a simplistic way) through our framework [55, 56, 57]. And for symmetric memory effects, the stability condition we derived offers a natural and precise generalization of the rule that environmental modification must produce negative feedbacks for every species.

Aside from clarifying and generalizing the conditions for coexistence, our model makes several distinct predictions about the relationship between diversity and robustness. When memory effects are symmetric, more diverse communities can tolerate higher local extinction rates. This phenomenon is due to “dilution effects” closely related to those studied in disease ecology [58]. If memory effects decay in time, the benefits of diversity may be even greater. In the bistable regime, high diversity in terms of species richness *and* relative abundance is needed to maintain coexistence. However, a negative relationship between diversity and robustness is observed when communities form a successional cycle, indicating that varied behaviors are possible even in this simple model, and that qualitatively distinct rate structures can support coexistence.

In the cases where diversity promotes robustness, coexistence may be sensitive to species loss as a result. Fig. 2C shows an example where the removal of a single species causes the extinction of the entire community. In the waning memory model, even perturbing some species to low abundance may cause such a collapse. These outcomes are most likely in marginal environments (high *m*). The dependence of long-term outcomes on the magnitude of *m*—observed in the nonsymmetric case, as well, through the loss of stability as *m* increases— is a surprising behavior that has no analog in simple models of community dynamics that do not incorporate local environmental feedbacks (e.g., generalized Lotka-Volterra models).

Our model is not meant to capture the detailed behavior of any particular community, but instead to highlight and investigate the essential dynamics common to a range of systems. This approach strikes a balance between models that explicitly incorporate the mechanisms of environmental modification [5, 59, 60] and those that fold environmental modification into direct interactions between species [2, 13, 14]. We view our approach as complementary to both. Despite its simplicity, our model exhibits rich and informative dynamics. It may also be possible to parameterize and test our model in empirical systems where investigators have measured parameters akin to our colonization rate matrices [4, 7, 8].

Inevitably, this approach requires us to make several strong assumptions about the underlying metapopulation dynamics of the community. For instance, we ignore the possibility of age or stage structures; complex spatial structures; co-occupancy of patches; and direct interactions between species. Many or all of these factors will be present in natural landscapes, and each has been shown to affect the dynamics of model communities [31, 33, 37]. However, by omitting these and other factors known to shape community dynamics, we are able to isolate the behavior of environmentally-mediated interactions, especially in relation to species coexistence. Integrating additional processes will likely raise interesting questions. To take one example, strong spatial structure, which is a central feature of many of the systems discussed here, might stabilize oscillatory dynamics, like those shown in Fig. 4, by coupling out of sync patches [21] or distributing oscillations over space [61]. Alternatively, though, localized dispersal could lead to spatial segregation of species, decoupling the successional cycles or negative feedbacks that sustain community diversity.

We have shown that our main findings are robust to relaxing one central assumption of the model—the permanence of memory effects—although qualitatively new behaviors arise in this case, as well. Many natural systems likely violate our model assumptions in the opposite direction, by exhibiting memory effects that endure beyond each new colonization event. Adaptive immunity, where memory effects may be lifelong, is an obvious example [25, 27, 28], but many others are possible. It is straightforward to extend our model to include longer-term memory effects, but not to analyze such extensions, which require a precipitous expansion of the state space and model complexity [26, 27]. It may be possible to consider other approximations, such as memory influencing extinction rates, or patch memory of the last two or three occupants—although even characterizing the behavior of our simple model under the most general (nonsymmetric) parameterizations remains an open problem. Expanding ecological theory to account for environmental feedbacks—beyond resource competition—on multiple spatial and temporal scales remains a key challenge and opportunity for further investigation.

## Materials and Methods

All computations (simulations and calculation of eigenvalues) were conducted in R (version 3.6.3). Where dynamics are shown, the model equations (Eq. 1) were solved numerically using the deSolve package with the “ode45” Runge-Kutta method. Coexistence conditions shown in Fig. 5A were obtained using Mathematica (see SI Waning Memory Effects for details). Code to generate all figures and simulation results is available on GitHub [45].

## Acknowledgments

We thank C. Serván, A. Skwara, P. Lemos, and J. Grilli for helpful discussion. J. Bergelson, M.O. Carlson, P. Lechon, and two anonymous reviewers provided thoughtful feedback that improved the text.

