## Supplementary material for "Metapopulations with habitat modification": SI Appendix

#### 1 Model Equations and Coexistence Equilibrium

##### 1.1 Model equations

It is convenient to express our model (Eq. 1 in the main text) in matrix form:

$$\begin{aligned}\frac{d\boldsymbol{x}(t)}{dt} &= D(\boldsymbol{x}(t))\left(-\boldsymbol{m} + P\boldsymbol{y}(t)\right)\\ \frac{d\boldsymbol{y}(t)}{dt} &= \left(D(\boldsymbol{m}) - D(\boldsymbol{y}(t))P^T\right)\boldsymbol{x}(t)\end{aligned} \tag{1}$$

Here, $D(\boldsymbol{z})$ denotes a diagonal matrix with vector $\boldsymbol{z}$ on the diagonal. Vectors $\boldsymbol{x}(t) \in \mathbb{R}^n$ and $\boldsymbol{y}(t) \in \mathbb{R}^n$ express the frequency (at time $t$) of patches occupied by species $i$ and vacant patches last occupied by species $i$ ("in state $i$"), respectively. $\boldsymbol{m} \in \mathbb{R}^n$ is a vector of local extinction rates, $m_1, \ldots, m_n > 0$, and $P \in \mathbb{R}^{n\times n}$ is a matrix of non-negative colonization rates.

As discussed in the main text, this system of equations generalizes Levins's classical metapopulation model in two ways: (i) there are $n$ species in the landscape, which interact by competing for patches, and (ii) vacant patches retain a "memory" of their last resident species, which determines the rate of re-colonization by every species in the community. The fact that each patch may be occupied by only a single species at any time (and therefore species compete for free patches) means that every patch is counted exactly once in the state variables $\boldsymbol{x}(t)$ and $\boldsymbol{y}(t)$. This implies zero-sum dynamics, as verified by summing:

$$\sum_i \frac{dx_i(t)}{dt} + \frac{dy_i(t)}{dt} = \sum_i \left(-m_i\, x_i(t) + x_i(t)\sum_{j=1}^n p_{ij}\, y_j(t) + m_i\, x_i(t) - y_i(t)\sum_{j=1}^n p_{ji}\, x_j(t)\right) = 0\,.$$

Patches, therefore, are never created or destroyed through the dynamics. Since the invariant quantity $T = \sum_{i=1}^n x_i + \sum_{i=1}^n y_i$ is arbitrarily determined by the initial conditions, we assume throughout that $T = 1$; in other words, $\boldsymbol{x}(t)$ and $\boldsymbol{y}(t)$ are frequencies.

In the following sections, we analyze the model (Eq. 1) in detail for arbitrary $n$ and considering a range of assumptions. On page 10, we include an analysis of the case $n = 2$, which summarizes and illustrates much of this analysis as applied to a more concrete example.

#### 1.2 Coexistence equilibrium

The system defined by Eq. 1 admits $2^n$ equilibria, corresponding to distinct combinations of species presence/absence (and counting the degenerate equilibrium with $\boldsymbol{x}(t) = 0$). In this study, we are primarily interested in the existence and stability of the unique equilibrium where all species (and patch states) are present at non-zero frequency – we refer to this as the *coexistence equilibrium.* To solve for the coexistence equilibrium (frequencies $\boldsymbol{x}^*$ and $\boldsymbol{y}^*$, dropping the time dependence), we first set $\frac{d\boldsymbol{x}}{dt} = 0$, which yields the linear system

$$P\boldsymbol{y}^* = \boldsymbol{m} \tag{2}$$

for $\boldsymbol{y}^*$. When $P$ has full rank, the equilibrium frequencies are found by matrix inversion: $\boldsymbol{y}^* = P^{-1}\boldsymbol{m}$. Invertibility of $P$ requires (at a minimum) that patch memory effects operate in the system. If not, then $p_{ij} = p_{ik} = p_i$ for all $i, j$, and $k$ (i.e., $P$ has constant rows), which means $P$ is a rank-one matrix, and therefore non-invertible for any $n > 1$. In this case, Eq. 2 admits a solution only if $\boldsymbol{m}$ is exactly proportional to the columns of $P$. If not, there is no equilibrium at all. However, even if this proportionality holds, there is no robust equilibrium. This situation, corresponding to a perfect trade-off in species' colonization and extinction rates, is degenerate, and unbiological for two reasons: (i) it requires fine-tuning of the parameters, and (ii) there are infinitely many equilibria, which means that all but one species will eventually drift to zero frequency (extinction) in a system of finite size. For the rest of the analysis, we assume that memory effects do operate, and that the colonization rate structure is non-degenerate, so $P$ is invertible.

Solving for the equilibrium species frequencies $\boldsymbol{x}^*$ is less straightforward. Substituting $\boldsymbol{y}^* = P^{-1}\boldsymbol{m}$ into $\frac{d\boldsymbol{y}}{dt}$ gives us the system

$$\left(D(\boldsymbol{m}) - D(P^{-1}\boldsymbol{m})P^T\right)\boldsymbol{x}^* = 0. \tag{3}$$

Let us define $Q = D(\boldsymbol{m}) - D(P^{-1}\boldsymbol{m})P^T$. The equilibrium frequencies $\boldsymbol{x}^*$ correspond to the one-dimensional null space of $Q$, or equivalently, an eigenvector of $Q$ with eigenvector 0. In general, then, there is no closed form expression for $\boldsymbol{x}^*$ (although these frequencies are easily computed numerically). However, for the important purpose of determining whether the coexistence equilibrium is biologically feasible (having all positive components), it is not necessary to compute $\boldsymbol{x}^*$ – only $\boldsymbol{y}^*$ is needed. This is because the matrix $-Q^T$ has the special structure of a transition rate matrix, which means that it satisfies:

1. $-q_{ii} > 0$ for all $i$
2. $-q_{ij} < 0$ for all $i \neq j$
3. $\sum_i q_{ij} = 0$ for all $j$ .

All three conditions are easily verified by observing that $Q = D(P\boldsymbol{y}^*) - D(\boldsymbol{y}^*)P^T$ (using the equilibrium relationship $\boldsymbol{m} = P\boldsymbol{y}^*$). Every transition rate matrix possesses a zero eigenvalue corresponding to an eigenvector of constant sign (the stationary distribution of the corresponding continuous-time Markov chain). This establishes that either all $x_i > 0$ or all $x_i < 0$. Eq. 3 determines $\boldsymbol{x}^*$ only up to a multiplicative constant; in order to determine the constant (and thus the sign), we invoke the fact that $1 = \sum_{i=1}^n x_i + \sum_{i=1}^n y_i$. Let $C$ be the sum of $\boldsymbol{x}^*$ at equilibrium. Then we have

$$\begin{aligned} 1 &= \mathbf{1}^T(\boldsymbol{x} + \boldsymbol{y}) \\ &= C + \mathbf{1}^T P^{-1}\boldsymbol{m} \end{aligned} \tag{4}$$

and so

$$C = 1 - \mathbf{1}^T P^{-1}\boldsymbol{m} \tag{5}$$

where $\mathbf{1}^T$ is a vector of $n$ ones. Combining these facts, all components of $\boldsymbol{x}^*$ are positive only if $\mathbf{1}^T P^{-1}\boldsymbol{m} < 1$; in other words, only if the vacant patch frequencies sum to less than 1.

The coexistence equilibrium is therefore feasible if and only if both $P^{-1}\boldsymbol{m} > 0$, elementwise, and $\mathbf{1}^T P^{-1}\boldsymbol{m} < 1$. Clearly these two criteria may be checked without reference to $\boldsymbol{x}^*$.

##### 1.3 Equilibrium species frequencies in special cases

Under various conditions, it becomes possible to express the equilibrium species frequencies in a straightforward way. Most notably, when $P$ is symmetric (i.e., $P = P^T$), the species frequencies $\boldsymbol{x}^*$ are proportional to the vacant patch frequencies $\boldsymbol{y}^*$. This is verified by setting $\boldsymbol{x}^* = k\boldsymbol{y}^*$, where $k$ is a constant of proportionality, and then substituting into Eq. 3:

$$\begin{aligned} \left(D(\boldsymbol{m}) - D(P^{-1}\boldsymbol{m})P^T\right)\boldsymbol{x} &= k(D(\boldsymbol{m}) - D(P^{-1}\boldsymbol{m})P)P^{-1}\boldsymbol{m} \\ &= k(D(\boldsymbol{m})P^{-1}\boldsymbol{m} - D(P^{-1}\boldsymbol{m})\boldsymbol{m}) \\ &= 0. \end{aligned} \tag{6}$$

As when assessing feasibility, the constant $k$ must be determined using the zero-sum constraint. We find

$$k = \frac{1}{\sum \boldsymbol{y}^*} - 1 = \frac{1}{\mathbf{1}^T P^{-1}\boldsymbol{m}} - 1. \tag{7}$$

It is also possible to write $\boldsymbol{x}^*$ directly when $P$ is symmetrizable, meaning $P$ is of the form $DS$, where $D$ is a diagonal matrix and $S$ is symmetric. Then, we find $\boldsymbol{y}^* = S^{-1}D^{-1}\boldsymbol{m}$ and, following a calculation similar to Eq. 6, $\boldsymbol{x}^* = kD^{-1}\boldsymbol{y}^* = k(DSD)^{-1}\boldsymbol{m}$, with $k$ once again a constant to be determined.

An explicit expression for $\boldsymbol{x}^*$ is possible in other circumstances, as well, but these two cases will be needed in the following sections.

#### 2 Species-Specific Memory Effects

##### 2.1 Simplest case

We begin by studying the simplest non-trivial parameterization of the model. Let $P = D(\boldsymbol{\alpha}) + \beta J$, where $\boldsymbol{\alpha} = (\alpha_1, \ldots, \alpha_n)^T$ is a vector of species-specific memory effects (of any sign) and $\beta > 0$ expresses the background colonization rate for all species. We must have $\alpha_i \geq -\beta$ to ensure all rates are non-negative. Here, $J$ denotes the $n \times n$ matrix of ones, $J = \mathbf{1}\mathbf{1}^T$. Initially, we will assume that all species experience identical local extinction rates, $m$.

Under these assumptions, $P$ is a rank-one perturbation of a diagonal matrix, and it is possible to compute $P^{-1}$ using the Sherman-Morrison formula [1]. We find that the equilibrium frequencies are

$$y_i^* = \frac{1}{\alpha_i} \left( \frac{m}{1 + \beta\, R(\boldsymbol{\alpha})} \right) \tag{8}$$

where $R(\boldsymbol{\alpha})$ is the sum of reciprocals of the $\alpha_i$, $R(\boldsymbol{\alpha}) = \sum_i \frac{1}{\alpha_i}$. Using Eqs. 6 and 7, we also have

$$x_i^* = \frac{1}{\alpha_i} \left( \frac{R(\boldsymbol{\alpha})^{-1} + \beta - m}{1 + \beta\, R(\boldsymbol{\alpha})} \right). \tag{9}$$

In Eq. 9, the factor in parentheses has no dependence on $i$. This shows that the equilibrium frequency of each species is inversely proportional to the strength of its memory effects. This also makes clear that the equilibrium can only be feasible if all $\alpha_i$ share the same sign. We will assume this is case; otherwise, coexistence of all species at equilibrium is not possible, and some species will go extinct.

##### 2.2 Global stability

For this equilibrium to be an attractor of the system, it must be stable as well as feasible. We would ideally like to know whether the equilibrium is *globally stable*, meaning the dynamics approach the equilibrium from any initial condition (assuming all species are initially present at some non-zero frequency). Global stability can be established by constructing a Lyapunov function for the dynamics. A Lyapunov function for an autonomous dynamical system $\frac{d\boldsymbol{z}}{dt} = g(\boldsymbol{z})$, with $\boldsymbol{z} \in \mathbb{R}^n$, is a continuous scalar function $V : \mathbb{R}^n \to \mathbb{R}$ that has continuous first derivatives and satisfies the following:

1. $V(\boldsymbol{z}) > 0$ except for an equilibrium point $\boldsymbol{z}^*$, where $V(\boldsymbol{z}^*) = 0$
2. $\frac{dV}{dt} < 0$ except when $\boldsymbol{z} = \boldsymbol{z}^*$, where $\frac{dV}{dt} = 0$

In words, $V$ is a positive function that is strictly decreasing through the dynamics, until reaching a minimum at $\boldsymbol{z}^*$. The existence of such a function implies that the point $\boldsymbol{z}^*$ is globally Lyapunov stable. For a thorough discussion of Lyapunov functions and global stability, see [2, 3].

Unfortunately, there is no general method to construct Lyapunov functions. For some well-studied models in ecology, including MacArthur's consumer-resource model [4] and the generalized Lotka-Volterra (GLV) model [5], candidate Lyapunov functions are known, and can be used to show global stability under appropriate parameterizations. While our model is not among these, we take advantage of an embedding technique from dynamical systems theory to make use of the known results for GLV.

Brenig [6] has shown that GLV is a canonical model, in the sense that any quasi-polynomial (QP) system of ordinary differential equations can be recast into GLV form through a change of variables. QP systems take the form

$$\frac{du_i}{dt} = u_i \left( \lambda_i + \sum_{j=1}^{m} A_{ij} \prod_{k=1}^{l} u_k^{B_{jk}} \right), \quad i = 1, ..., l \tag{10}$$

where $A$ is an $l \times m$ matrix and $B$ is an $m \times l$ matrix, with $m \geq l$. Many ecological models are in QP form, including ours. By casting our model from QP into GLV form, it becomes possible to use a class of candidate Lyapunov functions for GLV to prove global stability. Typically, this process requires an enlargement of the state space, and therefore the embedded dynamics are subject to constrained initial conditions. As we will show, these constraints are crucial for our stability analysis.

More concretely, our model is a QP system with $\boldsymbol{u}(t) = (\boldsymbol{x}(t), \boldsymbol{y}(t))^T$, $\boldsymbol{\lambda} = (-\boldsymbol{m}, 0)^T$, and $A$ and $B$ are block-structured matrices

$$A = \begin{pmatrix} 0 & P & 0 \\ -P^T & 0 & mI \end{pmatrix} \quad B = \begin{pmatrix} I & 0 \\ 0 & I \\ I & -I \end{pmatrix} \tag{11}$$

where each block is $n \times n$ (so $l = 2n$ and $m = 3n$). Such a system can be recast into GLV form with state variables

$$z_i = \prod_{j=1}^{m} u_j^{B_{ij}}, \quad i = 1, ..., m \tag{12}$$

and dynamics given by

$$\frac{d\boldsymbol{z}}{dt} = D(\boldsymbol{z})(B\boldsymbol{\lambda} + BA\boldsymbol{z}). \tag{13}$$

For more details, see [6, 7, 8]. In our particular case, this procedure amounts to defining the new variables $r_i = \frac{x_i}{y_i}$, with the resulting $3n$-"species" GLV system defined by variables $\boldsymbol{z}(t) = (\boldsymbol{x}(t), \boldsymbol{y}(t), \boldsymbol{r}(t))^T$, growth rates $B\boldsymbol{\lambda} = (-\boldsymbol{m}, 0, -\boldsymbol{m})^T$, and interaction matrix

$$BA = \begin{pmatrix} 0 & P & 0 \\ -P^T & 0 & mI \\ P^T & P & -mI \end{pmatrix}. \tag{14}$$

The interaction matrix for this larger system is singular, and the models defined by Eq. 1 and Eqs. 13-14 are equivalent only on the manifold where $r_i = \frac{x_i}{y_i}$ for all $i$.

All of these manipulations finally put us in a position to try applying a well-known family of candidate Lyapunov functions for GLV. The function

$$V(\boldsymbol{z}) = \sum_i \sigma_i \left( z_i - z_i^* - z_i^* \log\left(\frac{z_i}{z_i^*}\right)\right), \tag{15}$$

with non-negative constants $\sigma_i$, is smooth and always positive except at $V(\boldsymbol{z}^*) = 0$. Thus, $V$ is a Lyapunov function for the GLV system with equilibrium $\boldsymbol{z}^*$ if there is a choice of $\boldsymbol{\sigma}$ such that $\frac{dV}{dt} < 0$ for all $\boldsymbol{z} \neq \boldsymbol{z}^*$ [5]. The quantity $\frac{dV}{dt}$ conveniently reduces to

$$(\boldsymbol{z} - \boldsymbol{z}^*)^T \left(\frac{D(\boldsymbol{\sigma})M + M^T D(\boldsymbol{\sigma})}{2}\right)(\boldsymbol{z} - \boldsymbol{z}^*) \tag{16}$$

where $M$ is the interaction matrix (in our case, $M = BA$).

We can now show that the equilibrium found in Eqs. 8-9 is globally stable for our simplest model. Let $\boldsymbol{\sigma} = (\mathbf{1}, \mathbf{1}, \frac{-1}{k+1} D(\boldsymbol{\alpha})^{-1}\mathbf{1})^T$, where $k$ is the constant of proportionality $\frac{R(\boldsymbol{\alpha})+\beta-m}{m}$. We use the short-hand $\Delta\boldsymbol{x} = \boldsymbol{x} - \boldsymbol{x}^*$, and similarly for $\boldsymbol{y}$ and $\boldsymbol{r}$. Evaluating Eq. 16, and noting the symmetry of $P$, yields

$$\frac{dV}{dt} = m\Delta\boldsymbol{y}^T\Delta\boldsymbol{r} - \frac{m}{k+1}(\Delta\boldsymbol{x} + \Delta\boldsymbol{y})^T P D(\boldsymbol{\alpha})^{-1}\Delta\boldsymbol{r} + \frac{m^2}{k+1}\Delta\boldsymbol{r}^T D(\boldsymbol{\alpha})^{-1}\Delta\boldsymbol{r} \tag{17}$$

$$= m\Delta\boldsymbol{y}^T\Delta\boldsymbol{r} - \frac{m}{k+1}(\Delta\boldsymbol{x} + \Delta\boldsymbol{y})^T (D(\boldsymbol{\alpha}) + \beta J) D(\boldsymbol{\alpha})^{-1}\Delta\boldsymbol{r} + \frac{m^2}{k+1}\Delta\boldsymbol{r}^T D(\boldsymbol{\alpha})^{-1}\Delta\boldsymbol{r} \tag{18}$$

which simplifies to

$$= m\Delta\boldsymbol{y}^T\Delta\boldsymbol{r} - \frac{m}{k+1}(\Delta\boldsymbol{x} + \Delta\boldsymbol{y})^T\Delta\boldsymbol{r} + \frac{m^2}{k+1}\Delta\boldsymbol{r}^T D(\boldsymbol{\alpha})^{-1}\Delta\boldsymbol{r} \tag{19}$$

$$= \frac{-m}{k+1}(\Delta\boldsymbol{x} - k\Delta\boldsymbol{y})^T\Delta\boldsymbol{r} + \frac{m^2}{k+1}\Delta\boldsymbol{r}^T D(\boldsymbol{\alpha})^{-1}\Delta\boldsymbol{r} \tag{20}$$

using the zero-sum constraint, $(\Delta\boldsymbol{x} + \Delta\boldsymbol{y})^T J = 0$. Next we recall that $r_i = \frac{x_i}{y_i}$, which, along with the proportionality of $\boldsymbol{y}^*$ and $\boldsymbol{x}^*$, implies that $\Delta\boldsymbol{r} = D(\boldsymbol{y})^{-1}(\Delta\boldsymbol{x} - k\Delta\boldsymbol{y})$, and so

$$\frac{dV}{dt} = \frac{1}{k+1}(-m\Delta\boldsymbol{r}^T D(\boldsymbol{y})\Delta\boldsymbol{r} + m^2\Delta\boldsymbol{r}^T D(\boldsymbol{\alpha})^{-1}\Delta\boldsymbol{r}). \tag{21}$$

Because the frequencies $\boldsymbol{y}$ are always non-negative, and $\Delta\boldsymbol{r}^T D(\boldsymbol{y})\Delta\boldsymbol{r}$ is a quadratic form, the first term is always negative. The second term is always negative if $D(\boldsymbol{\alpha})^{-1}$ is a negative definite matrix. This requires $\alpha_i < 0$ for all $i$. These calculations show that when all species have negative memory effects, the coexistence equilibrium is globally stable. It is straightforward to show using similar arguments that if all $\alpha_i > 0$, the coexistence equilibrium is never stable, except when $n = 1$. In other words, when species exhibit positive memory effects, the only long-term outcome is monodominance.

While our Lyapunov function can be written entirely in terms of the original model variables and parameters as

$$
\begin{aligned}
V &= \sum_i x_i - x_i^* - x_i^* \log\left(\frac{x_i}{x_i^*}\right) + y_i - y_i^* - y_i^* \log\left(\frac{y_i}{y_i^*}\right) - \frac{1}{(k+1)\alpha_i}\left(\frac{x_i}{y_i} - k - k\log\left(\frac{x_i}{ky_i}\right)\right) \\
&= \sum_i -x_i^* \log\left(\frac{x_i}{x_i^*}\right) - y_i^* \log\left(\frac{y_i}{y_i^*}\right) - \frac{1}{(k+1)\alpha_i}\left(\frac{x_i}{y_i} - k - k\log\left(\frac{x_i}{ky_i}\right)\right)
\end{aligned}
\tag{22}
$$

and verified without reference to the GLV embedding, we trace the process of recasting in GLV form to demonstrate the utility of this approach. The difficulty of generating suitable Lyapunov functions – which is usually seen a matter of inspired guesswork or laborious trial-and-error – is a major obstacle to their wider application. The canonical status of GLV, and the potential to exploit this fact for constructing Lyapunov functions, has been known for decades, but very rarely utilized in ecology (but see [9]). Our derivation illustrates how GLV embedding can systematize the search for a Lyapunov function, by reducing the problem to a choice of appropriate constants (here, $\sigma_i$). In many cases, this is likely to be a more fruitful effort than divining an appropriate function directly.

##### 2.3 Feasibility and diversity

We now return to consideration of feasibility. We have seen that feasibility requires that all $\alpha_i$ share a common sign, and stability requires that all $\alpha_i < 0$. When these conditions are satisfied, what else is needed to guarantee feasibility? The remaining criterion is a community-wide condition:

$$m < R(\boldsymbol{\alpha})^{-1} + \beta. \tag{23}$$

When this inequality holds, and assuming all $\alpha_i < 0$, both $\boldsymbol{x}^*$ and $\boldsymbol{y}^*$ are strictly positive. Eq. 23 gives an upper bound on the local extinction rate that the coexisting community can tolerate. Because $\alpha_i < 0$, the quantity $R(\boldsymbol{\alpha})^{-1}$ is always negative, but it increases toward zero as any new species is added to the community. As a consequence, the maximum $m$ compatible with feasibility, $m_{\max}$, increases with diversity, as well. This implies a positive diversity-robustness relationship: Assuming some environmental component to $m$ (habitat quality, external disturbance rate, etc.), more speciose communities can tolerate environments that would drive a subset of the species to extinction. In this scheme, species with weaker memory effects (i.e. $\alpha_i$ closer to 0) contribute more strongly to raising $m_{\max}$. Intuitively, these species are more abundant at equilibrium, and more effectively dilute the negative memory effects of other species. As species join the community, $m_{\max}$ increases to an asymptotic upper bound at $m_{\max} = \beta$. In this limit, the memory effects in the system are so diluted that the effective colonization rate for all species is just $\beta$, the background rate.

It is indicative of the contrasting effects of diversity in this model that species with weak negative memory effects provide a strong community benefit (higher $m_{\max}$) but also occupy a large fraction of patches. Coexisting communities blur the distinction between mutualism and competition, as species benefit one another through a dilution effect, but compete for available patches. One consequence is a non-monotonic relationship between diversity and equilibrium frequency. Consider the equilibrium value, $x_i^*$, for a focal species as the community composition varies with $m$ constant. Adding or removing species affects $x_i^*$ through the quantity $R(\boldsymbol{\alpha})$, which we take for a moment as a continuous variable, $R$, that ranges from $\frac{1}{m-\beta}$ (species $i$ alone with strong negative memory effects) to $-\infty$ (very many species, or species with very weak negative memory effects). The derivative of $x_i^*$ with respect to $R$ is

$$\frac{dx_i^*}{dR} = \frac{-1}{\alpha_i}\left(\frac{1}{R^2} - \frac{m\beta}{(1+\beta R)^2}\right) \tag{24}$$

which changes sign as $R$ tends toward $-\infty$. For $R \approx \frac{1}{m-\beta}$, recalling that $\alpha_i < 0$, we have $\frac{dx_i^*}{dR} > 0$, indicating that $x_i^*$ increases with increasing diversity (decreasing $R$). However, as $R$ becomes very negative, $\frac{dx_i^*}{dR}$ approaches 0 from above. The sign of $\frac{dx_i^*}{dR}$ changes once at

$$R = \frac{1}{m-\beta}\left(1 + \sqrt{\frac{m}{\beta}}\right) \tag{25}$$

which can be found by setting Eq. 24 equal to 0 (the other root always falls above $\frac{1}{m-\beta}$). Once $R$ drops below the critical value in Eq. 25, any increase in diversity decreases $R$ further, decreasing $x_i^*$, as well.

##### 2.4 Variation in local extinction rates

How does this picture change when species differ in their local extinction rates? Now, we allow each species to have a distinct local extinction rate, $m_i > 0$. It turns out that stability is unaffected by this variation, although feasibility may be. And while the relationship between diversity and robustness becomes more complex in this case, we will see that there is still a well-defined positive relationship between the two.

Using the Sherman-Morrison formula, as before, we find

$$y_i^* = \frac{1}{\alpha_i}\left(m_i - \frac{\beta(\sum_j \frac{m_j}{\alpha_j})}{1+\beta R(\boldsymbol{\alpha})}\right). \tag{26}$$

Using Eq. 7, we also have

$$x_i^* = \frac{1}{\alpha_i}\left(m_i - \frac{\beta(\sum_j \frac{m_j}{\alpha_j})}{1+\beta R(\boldsymbol{\alpha})}\right)\left(\frac{1+\beta R(\boldsymbol{\alpha})}{\sum_j \frac{m_j}{\alpha_j}} - 1\right). \tag{27}$$

As before, all $\alpha_i$ must share the same sign for feasibility. For stability, this common sign must be negative. This can be verified using a Lyapunov function of the same form as Eq. 15, but with $\boldsymbol{\sigma} = \{\mathbf{1}, \mathbf{1}, \frac{-1}{k+1}D(\boldsymbol{m})D(\boldsymbol{\alpha})^{-1}\mathbf{1}\}$.

The requirement that $\alpha_i < 0$ induces a positive diversity-robustness relationship in this case, as well. The picture is complicated by the fact that each species may react differently to a set of environments – a favorable environment for one species (decreased $m_i$) may be unfavorable for another (increased $m_j$). However, any community-environment pair can be characterized by a weighted average of of the local extinction rates: $w(\boldsymbol{m}) = \sum_j \left(\frac{R(\boldsymbol{\alpha})}{\alpha_j}\right) m_j$. Then a community-wide feasibility condition, directly analogous to Eq. 23, is

$$w(\boldsymbol{m}) < R(\boldsymbol{\alpha})^{-1} + \beta\,. \tag{28}$$

The interpretation of Eq. 28 is nearly identical to Eq. 23: $R(\boldsymbol{\alpha})^{-1}$ is negative and increasing with species richness, so diverse communities are able to tolerate higher $w(\boldsymbol{m})$.

Unlike the constant $m$ case, Eq. 28 is not sufficient for feasibility. We also find a requirement that no particular $m_i$ is too much larger than the rest. From Eq. 27, one can show that the inequality

$$\frac{m_i}{w(\boldsymbol{m})} < \frac{\beta R(\boldsymbol{\alpha})}{1 + \beta R(\boldsymbol{\alpha})} \tag{29}$$

must hold for all $i$. This feasibility condition sets an upper bound on the ratio of each species' local extinction rate to the weighted average of the whole community. When diversity is low, this condition is not very restrictive, but as diversity increases and $R(\boldsymbol{\alpha})$ decreases toward $-\infty$, the allowable variation in $m_i$ becomes very small.

### 3 Symmetric Memory Effects

In this section we consider arbitrary symmetric memory effects (i.e., $P = P^T$). For two species, every symmetric matrix has constant off-diagonals, so the results of the previous section apply. For $n > 2$, we derive a stability condition that naturally, but precisely, generalizes the intuitive notion that all species must have a disadvantage colonizing "their own" vacant patches (patches of type $i$, for species $i$), compared to vacant patches in other states.

As $n$ grows, it quickly becomes impractical to write the equilibrium frequencies explicitly. The general feasibility conditions in SI Model Equations and Coexistence Equilibrium must be checked in each case. For the remainder of this section we assume the coexistence equilibrium is feasible and focus on its stability properties. In subsection Feasibility and Diversity, we return to consider the diversity-robustness relationship, as suggested by Eqs. 23 and 28.

#### 3.1 Local stability

Allowing $P$ to be an arbitrary (non-negative) symmetric matrix makes the problem of constructing a Lyapunov function much more difficult. Instead, we consider the *local* stability of the coexistence equilibrium. Once again, we begin by assuming $m$ is identical for all species. However, in subsection Relaxing Assumptions we present numerical evidence that any locally stable equilibrium is in fact globally stable, and that variation in local extinction rates never affects stability.

Local stability of the coexistence equilibrium depends on the real parts of the eigenvalues of the Jacobian matrix, $J^*$, evaluated at $(\boldsymbol{x}^*, \boldsymbol{y}^*)^T$. Due to the zero-sum constraint, the model dynamics are confined to the simplex, and there is necessarily one zero eigenvalue, which corresponds to an eigenvalue pointing "out" of the simplex. This eigenvalue has no bearing on stability. The coexistence equilibrium is then locally asymptotically stable if and only if the remaining $2n - 1$ eigenvalues of $J^*$ all have negative real part. In general, we have the Jacobian

$$J^* = \begin{pmatrix} 0 & D(\boldsymbol{x}^*)P \\ D(\boldsymbol{m}) - D(\boldsymbol{y}^*)P^T & -D(P^T\boldsymbol{x}^*) \end{pmatrix} \tag{30}$$

or, using the equilibrium relationship $D(\boldsymbol{m})\boldsymbol{x}^* = D(\boldsymbol{y}^*)P^T\boldsymbol{x}^*$,

$$J^* = \begin{pmatrix} 0 & D(\boldsymbol{x}^*)P \\ D(\boldsymbol{m}) - D(\boldsymbol{y}^*)P^T & -D(\boldsymbol{y}^*)^{-1}D(\boldsymbol{m})D(\boldsymbol{x}^*) \end{pmatrix} \tag{31}$$

which simplifies considerably to

$$J^* = \begin{pmatrix} 0 & kD(\boldsymbol{y}^*)P \\ mI - D(\boldsymbol{y}^*)P & -kmI \end{pmatrix} \tag{32}$$

under the assumptions $P = P^T$ and $m_i = m$.

It is actually more convenient to work with another matrix, $J'$, which is *similar* to $J^*$. As such, $J^*$ and $J'$ share the same eigenvalues. Let us define the change of basis matrix

$$U = \begin{pmatrix} D(\boldsymbol{y}^*)^{1/2} & 0 \\ 0 & D(\boldsymbol{y}^*)^{1/2} \end{pmatrix} \tag{33}$$

Then $J'$ is given by

$$\begin{aligned} J' &= U^{-1}J^*U \\ &= \begin{pmatrix} 0 & kD(\boldsymbol{y}^*)^{1/2}PD(\boldsymbol{y}^*)^{1/2} \\ mI - D(\boldsymbol{y}^*)^{1/2}PD(\boldsymbol{y}^*)^{1/2} & -kmI \end{pmatrix} \end{aligned} \tag{34}$$

For simplicity we introduce the shorthand $S = D(\boldsymbol{y}^*)^{1/2}PD(\boldsymbol{y}^*)^{1/2}$. Now we will see that the eigenvalues of $J'$ can be written in terms of the eigenvalues of $S$. The $i$th eigenvalue of $J'$, denoted $\lambda_i$, satisfies

$$J' \begin{pmatrix} \boldsymbol{u}_i \\ \boldsymbol{v}_i \end{pmatrix} = \lambda_i \begin{pmatrix} \boldsymbol{u}_i \\ \boldsymbol{v}_i \end{pmatrix} \tag{35}$$

for the corresponding eigenvector $(\boldsymbol{u}_i, \boldsymbol{v}_i)^T$, with components $\boldsymbol{u}_i$ and $\boldsymbol{v}_i$ each of length $n$. This gives us the system of equations

$$\begin{aligned} kS\boldsymbol{v}_i &= \lambda_i\boldsymbol{u}_i \\ m\boldsymbol{u}_i - S\boldsymbol{u}_i - mk\boldsymbol{v}_i &= \lambda_i\boldsymbol{v}_i. \end{aligned} \tag{36}$$

We notice that Eq. 36 can satisfied by $\boldsymbol{u}_i = c\boldsymbol{v}_i$, for an undetermined constant $c$, whenever $\boldsymbol{v}_i$ is an eigenvector of $S$. Denote the corresponding eigenvalue of $S$ as $\lambda(S)_i$, which must equal $\frac{c}{k}\lambda_i$, according to the first equation. Assuming $\lambda_i \neq 0$, we can then re-write the second equation entirely in terms of $\boldsymbol{v}_i$:

$$mk\frac{\lambda(S)_i}{\lambda_i}\boldsymbol{v}_i - k\frac{\lambda(S)_i}{\lambda_i}S\boldsymbol{v}_i - mk\boldsymbol{v}_i = mk\frac{\lambda(S)_i}{\lambda_i}\boldsymbol{v}_i - k\frac{\lambda(S)_i^2}{\lambda_i}\boldsymbol{v}_i - mk\boldsymbol{v}_i = \lambda_i\boldsymbol{v}_i \tag{37}$$

which yields the scalar equation

$$\begin{aligned} 0 &= mk\frac{\lambda(S)_i}{\lambda_i} - k\frac{\lambda(S)_i^2}{\lambda_i} - mk - \lambda_i \\ &= \lambda_i^2 + km\lambda_i + k\lambda(S)_i(\lambda(S)_i - m). \end{aligned} \tag{38}$$

Eq. 38 has the solutions

$$\lambda_i = \frac{-km \pm \sqrt{(km)^2 - 4k\lambda(S)_i(\lambda(S)_i - m)}}{2}. \tag{39}$$

We can see that each eigenvalue of $S$ determines a pair of eigenvalues $J'$, which we write as $\lambda_i^+$ and $\lambda_i^-$. If the coexistence equilibrium is feasible, $-km$ will always be negative, and so $\lambda_i^-$ will have negative real part. The real part of $\lambda_i^+$ will be negative if $\lambda(S)_i < 0$ or $\lambda(S)_i > m$, and exactly zero when $\lambda(S)_i = 0$ or $m$. In fact, one eigenvalue of $S$ is equal to $m$, with corresponding eigenvector $D(\boldsymbol{y}^*)^{1/2}\mathbf{1}$. This is easy to verify:

$$\begin{aligned} SD(\boldsymbol{y}^*)^{1/2}\mathbf{1} &= D(\boldsymbol{y}^*)^{1/2}PD(\boldsymbol{y}^*)^{1/2}D(\boldsymbol{y}^*)^{1/2}\mathbf{1} \\ &= D(\boldsymbol{y}^*)^{1/2}PD(\boldsymbol{y}^*)\mathbf{1} \\ &= mD(\boldsymbol{y}^*)^{1/2}PP^{-1}\mathbf{1} \\ &= mD(\boldsymbol{y}^*)^{1/2}\mathbf{1}. \end{aligned} \tag{40}$$

This eigenpair generates the expected zero eigenvalue of $J'$.

The matrix $S$ and its eigenvector $D(\boldsymbol{y}^*)^{1/2}\mathbf{1}$ are both non-negative, so $m$ is the Perron eigenvalue of $S$ [1]. This implies that all other eigenvalues of $S$ are smaller than $m$ in magnitude. In particular, there is no $\lambda(S)_i > m$. This means that the coexistence equilibrium is stable if an only if $m$ is the sole positive eigenvalue of $S$. Because $S$ and $P$ are symmetric and *congruent*, Sylvester's law of inertia guarantees that they share the same number of positive, negative, and zero eigenvalues [1]. Thus, we can equivalently state that the coexistence equilibrium is stable if and only if $P$ has a single positive eigenvalue.

This kind of matrix (nonnegative, symmetric, with exactly one positive eigenvalue) has previously been associated with the stable maintenance of polymorphisms in models in population genetics (e.g. [10, 11, 12]). These and other studies [13, 14, 15] have produced several characterizations of this class of matrix, which allow us to develop some biological intuition for the stability condition (see main text).

#### 3.2 Relaxing assumptions

While the analysis above relies on the assumption $m_i = m$ and shows only local stability, numerical simulations suggest that the same stability condition holds regardless of $\boldsymbol{m}$ (assuming feasibility), and that this stability is global in character.

To examine the consequences of variation in $\boldsymbol{m}$, we drew random symmetric $P$ matrices with $n = 4$ and $p_{ij} = p_{ji} \sim U[0, 1]$, and numerically checked local stability for different choices of $\boldsymbol{m}$. To assess whether the stability criterion derived with constant $m$ remained necessary and sufficient for stability in this generalized setting, we collected 1000 randomly distributed $P$ matrices with exactly one positive eigenvalue (putatively stable) and 1000 matrices with additional positive eigenvalues (putatively unstable). For each matrix, we considered 1000 random $\boldsymbol{m}$ vectors. To avoid sampling many choices of $\boldsymbol{m}$ incompatible with feasibility, which would be computationally costly, we first sampled a vector $\boldsymbol{l}$ uniformly from the $n$-simplex, which we took to be proportional to $\boldsymbol{y}^*$ (and consequently $\boldsymbol{x}^*$). Then we computed $\boldsymbol{m}$ as $cP\boldsymbol{l}$ where $c$ is a scalar uniformly distributed between 0 and 1. Finally, the equilibrium frequencies were computed accordingly. This sampling strategy is commonly used to draw random parameters compatible with feasibility (see, for example, [16, 17, 18]). Across all $10^6$ combinations of $P$ and $\boldsymbol{m}$ for each stability category (stable or unstable), we found that the stability of the system with variation in $m_i$ was always correctly predicted by the stability of the associated system with no variation. This suggests that the condition *$P$ has exactly one positive eigenvalue* is necessary and sufficient for stability of any community with symmetric memory effects, regardless of the values of each species' local extinction rates.

To check whether local stability implies global stability in this model, we again drew random symmetric $P$ matrices with $n = 4$ and $p_{ij} = p_{ji} \sim U[0, 1]$. For these simulations, we only retained $P$ matrices with exactly one positive eigenvalue, and sampled until we collected 1000 such matrices. For each realization of $P$, we calculated $m_{\max}$ and set $m = \frac{1}{2} m_{\max}$ for all species. Then we numerically integrated the model dynamics from 100 random initial conditions, sampled uniformly from the $2n$-simplex. In each case, we integrated the dynamics for 5000 time steps, and then checked if the trajectory had converged to the coexistence equilibrium. If not, we integrated for another 5000 time steps, and repeated this procedure up to 10 times. For all $10^5$ combinations of $P$ and initial conditions, we observed convergence to the equilibrium. This consistency strongly suggests that any locally stable equilibrium is also globally stable in our (symmetric) model. It is unnecessary to check the converse, in this case, because local instability implies that trajectories will always eventually move away from the equilibrium, regardless of initial conditions.

R scripts implementing these simulations are available on GitHub [19].

##### 3.3 Feasibility and diversity

Motivated by our analysis of the model with species-specific memory effects, it is natural to ask whether this stability condition for symmetric memory effects also induces a positive diversity-robustness relationship. To make this question precise we maintain the assumption of identical $m$ for all species, and consider the notion of an *assembly sequence* [20]. An assembly sequence is a sequence of sets of species beginning with a single species, and where each set contains the preceding set along with one additional species. For instance, $\{i\}$, $\{i, j\}$, $\{i, j, k\}$, where $i, j$ and $k$ are species labels, is an assembly sequence of length 3. For any set of species there is an upper bound, $m_{\max}$, on $m$ for feasibility of the coexistence equilibrium. We ask whether, for any coexisting set of species $S$, $m_{\max}$ increases along any assembly sequence that ends at $S$.

For this question to be meaningful, each set of species along the assembly sequence should coexist (for suitable $m$). Even if the equilibrium frequencies for $S$ all share the same sign, there is no guarantee that this property will hold for a subset. Thus, we need to assume this property holds for each set along the assembly sequence. On the other hand, given (potential) feasibility, it *is* the case that, if the equilibrium for $S$ is stable, then every subset will also be stable [13]. This is a straightforward consequence of the *eigenvalue interlacing theorem for bordered matrices* [1], which we will rely on again to prove that $m_{\max}$ is increasing along any assembly sequence. For our purposes, the theorem states that, if $P^k$ is a $k \times k$ symmetric matrix and $P^{k-1}$ is the matrix obtained by deleting the last row and column of $P^k$, then their respective eigenvalues, $\lambda_i^k$ and $\lambda_i^{k-1}$, can be ordered as:

$$\lambda_1^k \geq \lambda_1^{k-1} \geq \lambda_2^k \geq \lambda_2^{k-1} \cdots \geq \lambda_{k-1}^{k-1} \geq \lambda_k^k. \tag{41}$$

Using Eq. 41, along with the Perron-Frobenius theorem [1], we see that if $P^k$ is a positive symmetric matrix with exactly one positive eigenvalue, then $P^{k-1}$ must be as well. For any set of species, $S$, we can always order the corresponding colonization rate matrix, $P$, so that a subset of interest is obtained by sequentially removing the last row and column of $P$; the stability of any subset follows inductively.

We will also use the identity [21]

$$q_{ii}^{k-1} = q_{ii}^k - \frac{q_{in}^k}{q_{ni}^k} q_{nn}^k \quad \forall i = 1, \ldots, k-1 \tag{42}$$

which relates the elements of inverse matrices $(P^{k-1})^{-1} = (q_{ij}^{k-1})$ and $(P^k)^{-1} = (q_{ij}^k)$.

We are now in a position to prove the statement that $m_{\max}$ increases along any assembly sequence terminating in a coexisting set of species, $S$. $m_{\max}$ can be computed from Eq. 7, which gives

$$m_{\max} = \frac{1}{\mathbf{1}^T P^{-1} \mathbf{1}}. \tag{43}$$

We see that $m_{\max}$ is a decreasing function of the quantity $\mathbf{1}^T P^{-1} \mathbf{1}$. Thus, for $S$ with $k$ species and the preceding subset with $k-1$ species, we have $m_{\max}^k \geq m_{\max}^{k-1}$ if and only if $\mathbf{1}^T (P^k)^{-1} \mathbf{1} \leq \mathbf{1}^T (P^{k-1})^{-1} \mathbf{1}$. To manipulate this second inequality, we introduce the matrix $(\tilde{P}^{k-1})^{-1}$, which is just $(P^{k-1})^{-1}$ with a row and column of zeros appended. Now both $(P^k)^{-1}$ and $(\tilde{P}^{k-1})^{-1}$ are $k \times k$, so we can subtract:

$$\mathbf{1}^T (P^k)^{-1} \mathbf{1} - \mathbf{1}^T (P^{k-1})^{-1} \mathbf{1} = \mathbf{1}^T \left( (P^k)^{-1} - (\tilde{P}^{k-1})^{-1} \right) \mathbf{1} = \mathbf{1}^T R \, \mathbf{1} \leq 0. \tag{44}$$

Using Eq. 42, the elements of $R$ are simply $(r_{ij}) = \frac{(q_{in}^k)^2}{q_{nn}^k}$. Clearly the sign of the quadratic form in Eq. 44 is determined by the sign of $q_{nn}^k$. If this sign is always negative (or zero), then the inequality holds. As we have noted that the rows and columns of $P$ can be re-ordered to make any species the $n$th, we now show that $q_{nn}^k \leq 0$ by proving that every diagonal element of $(P^k)^{-1}$, the inverse of a positive symmetric matrix with exactly one positive eigenvalue, is non-positive.

This can be done by induction. In Eq. 44, we have at least $k = 2$. Taking this as our base case and computing the inverse explicitly we find

$$(P^{k=2})^{-1} = \frac{1}{\det(P^{k=2})} \begin{pmatrix} p_{22} & -p_{12} \\ -p_{21} & p_{11} \end{pmatrix} \tag{45}$$

which must have non-positive diagonal elements because $P^{k=2}$ is non-negative and the determinant is strictly negative. Now, in the induction step, we prove that if $q_{ii}^{k-1} \leq 0$ for all $i = 1, \ldots k-1$, then $q_{ii}^k \leq 0$ for all $i = 1, \ldots k$, as well. Here we use a proof by contradiction. First assume that $q_{kk}^k > 0$. Then, using Eq. 42, we must have $q_{ii}^k \geq q_{ii}^{k-1}$ for all $i = 1, \ldots k-1$. This implies that $\mathrm{Tr}((P^k)^{-1}) > \mathrm{Tr}((P^{k-1})^{-1})$. But the eigenvalues of $(P^k)^{-1}$ and $(P^{k-1})^{-1}$, which are $\frac{1}{\lambda_i^k}$ and $\frac{1}{\lambda_i^{k-1}}$, respectively, obey

$$\frac{1}{\lambda_1^{k-1}} \geq \frac{1}{\lambda_1^k} \geq 0 \geq \frac{1}{\lambda_k^k} \geq \frac{1}{\lambda_{k-1}^{k-1}} \cdots \geq \frac{1}{\lambda_2^{k-1}} \geq \frac{1}{\lambda_2^k}. \tag{46}$$

using Eq. 41 and the fact that both matrices have exactly one positive eigenvalue. These inequalities imply that $\mathrm{Tr}((P^k)^{-1}) < \mathrm{Tr}((P^{k-1})^{-1})$, a contradiction. Thus, we conclude that $q^k_{kk} < 0$. In this case, Eq. 42 shows that $q^k_{ii} \leq q^{k-1}_{ii}$ for all $i = 1, \ldots k-1$. These inequalities establish that if $q^{k-1}_{ii} \leq 0$ for all $i = 1, \ldots k-1$, then $q^k_{ii} \leq 0$ for all $i = 1, \ldots k$, so altogether we have shown that $q^k_{ii} \leq 0$ for any $k \geq 2$ and $i$.

Combined with the results above, this ultimately establishes that $m^k_{\max} \geq m^{k-1}_{\max}$ at every step along a (coexisting) assembly sequence. In this setting, a species-rich community can always tolerate a higher local extinction rate than any subset of its species. Notice that, if $m$ is constant, these results also imply that the total fraction of occupied patches at equilibrium increases along any assembly sequence. This kind of positive diversity-productivity relationship is commonly observed in empirical studies [22, 23, 24]. In our model, the relationships between diversity-robustness and diversity-productivity are closely connected, because the robustness benefit of diversity arises from a positive "dilution effect" of a more productive community, as discussed in the main text.

While we have shown here that the conditions for coexistence generically induce a positive diversity-robustness relationship, we are not able to quantify the magnitude of this effect, as in Eqs. 23 and 28. To get a sense of the possibilities, we simulated 500 assembly sequences, each terminating in a distinct coexisting community of 5 species. The matrix $P$ for each community was drawn at random with $p_{ij} = p_{ji} \sim U[0,1]$ i.i.d. for simplicity. We discarded matrices that did not permit coexistence, and sampled until we obtained 500 that did. For every community, we calculated $m_{\max}$ first for species 1 alone, then species 1 and 2 together, and so on, up to the full 5 species. If any subset along the assembly sequence lacked a feasible equilibrium, the entire sequence was discarded. As a comparison, we also simulated 500 assembly sequences terminating in communities having equilibria that were potentially feasible (i.e., $P^{-1}\mathbf{1} > 0$, elementwise), but unstable. Our procedure for these communities was identical, except that matrices were checked for instability, rather than stability. The results of these numerical experiments are shown in Fig. S1. For stable communities, $m_{\max}$ always increases with richness, as expected, and grows more than twofold on average from 1-species to 5-species communities. For unstable communities, we never observed $m_{\max}$ increase consistently along an assembly sequence, and Fig. S1 shows no trend in the average change of $m_{\max}$ with richness. This comparison makes it clear that the positive diversity-robustness relationship seen in coexisting communities is a consequence of the stability condition, rather than the feasibility conditions.

**Example: Coexistence criteria for a two-species system**

To help illustrate the ideas and quantitative results of the previous sections, we analyze the particular case where only two species are present in the landscape. These species, 1 and 2, are characterized by local extinction rates $m_1$ and $m_2$, as well as the rates at which they are able to colonize patches last occupied by a conspecific ($p_{11}$, for species 1 colonizing a patch last occupied by species 1, and similarly for $p_{22}$), and the rates at which they are able to colonize patches last occupied by the other species ($p_{12}$, for species 1 colonizing a patch last occupied by species 2, and similarly for $p_{21}$). These rates $p_{ij}$ encode differences in colonization ability across patch states for each species.

For example, if species 1 promotes local recruitment of specialized pathogens, we expect $p_{11} < p_{12}$. Or, if species 2 improves patch conditions for both species by breaking down unavailable nutrients into a labile form, then $p_{12} > p_{11}$ and $p_{22} > p_{21}$. Together the $p_{ij}$ constitute a matrix of rates, $P$:

$$P = \begin{pmatrix} p_{11} & p_{12} \\ p_{21} & p_{22} \end{pmatrix} . \tag{47}$$

Using these parameters, the model equations (Eq. 1) become

$$\begin{aligned}
\frac{dx_1(t)}{dt} &= -m_1\, x_1(t) + x_1(t)\,(p_{11}\, y_1(t) + p_{12}\, y_2(t)) \\
\frac{dx_2(t)}{dt} &= -m_2\, x_2(t) + x_2(t)\,(p_{21}\, y_1(t) + p_{22}\, y_2(t)) \\
\frac{dy_1(t)}{dt} &= m_1\, x_1(t) - y_1(t)\,(p_{11}\, x_1(t) + p_{21}\, x_2(t)) \\
\frac{dy_2(t)}{dt} &= m_2\, x_2(t) - y_2(t)\,(p_{12}\, x_1(t) + p_{22}\, x_2(t)) \ .
\end{aligned} \tag{48}$$

To find the unique coexistence equilibrium for this system, we first solve for the vacant patch frequencies ($y$ variables) at equilibrium. These are obtained from the equations governing the $x$ variables. The rates of change $\frac{dx_1}{dt}$ and $\frac{dx_2}{dt}$ are zero when

$$\begin{aligned}
-m_1 + p_{11}\, y_1 + p_{12}\, y_2 &= 0 \\
-m_2 + p_{21}\, y_1 + p_{22}\, y_2 &= 0
\end{aligned} \tag{49}$$

(neglecting cases where one of $x_1$ or $x_2$ is zero, which correspond to one-species subsystems). This is a special case of Eq. 2. The solutions to Eqs. 49 are

$$\begin{aligned}
y_1^* &= \frac{p_{12}m_2 - p_{22}m_1}{p_{12}p_{21} - p_{11}p_{22}} \\
y_2^* &= \frac{p_{21}m_1 - p_{11}m_2}{p_{12}p_{21} - p_{11}p_{22}}
\end{aligned} \tag{50}$$

which can only correspond to a biologically feasible equilibrium if both $y_1^*$ and $y_2^*$ are positive. To find the equilibrium frequencies for occupied patches, one can similarly obtain a pair of equations by setting $\frac{dy_1}{dt}$ and $\frac{dy_2}{dt}$ to zero and substituting in the $y^*$ values:

$$\begin{aligned}
(m_1 - p_{11}\, y_1^*)x_1 + p_{12}\, y_1^*\, x_2 &= 0 \\
(m_2 - p_{22}\, y_2^*)x_2 + p_{21}\, y_2^*\, x_1 &= 0\,.
\end{aligned} \tag{51}$$

However, using Eq. 49, both equations simplify to the identical relation

$$p_{12}\, y_2^*\, x_1 = p_{21}\, y_1^*\, x_2 \tag{52}$$

which has infinitely many solutions. This degeneracy reflects the fact that the homogeneous linear system of equations defining the $x^*$ is always singular, regardless of the choice of $P$ and $m$ (see Eq. 3 and accompanying text). Another way to express these solutions is

$$\begin{aligned} x_1^* &= \frac{k}{p_{12}} y_1^* \\ x_2^* &= \frac{k}{p_{21}} y_2^* , \end{aligned} \tag{53}$$

where $k$ is any constant. The particular $k$ defining equilibrium frequencies for the model is found by considering the fact that $x_1 + x_2 + y_1 + y_2$ must always equal 1. One obtains

$$k = \frac{p_{12}\, p_{21}\, (1 - y_1^* - y_2^*)}{p_{21}\, y_1^* + p_{12}\, y_2^*} . \tag{54}$$

From this expression and the equations above, it is clear that the $x^*$ share a common sign, which is positive whenever $1 > y_1^* + y_2^*$. This is the third and final requirement for the coexistence equilibrium to be feasible.

However, in order for these species to coexist at the equilibrium, it must also be (at least) locally stable. Even for this minimal example with only two species, directly analyzing the local stability with arbitrary parameters is extremely difficult. The difficulty arises because the system is essentially three-dimensional, not two, preventing the derivation of a simple formula for the eigenvalues of the Jacobian matrix. As in SI Symmetric Memory Effects (3.A), we must restrict our focus to the case where $m_1 = m_2$ and $p_{12} = p_{21}$. Even in this setting, computing the eigenvalues explicitly is a challenging task; instead, we can take advantage of the earlier results to see that the coexistence equilibrium is locally stable whenever $P$ has one positive and one negative eigenvalue. The eigenvalues of $P$ are

$$\lambda_1 = \frac{p_{11} + p_{22} + \sqrt{(p_{11} + p_{22})^2 - 4(p_{11}p_{22} - p_{12}^2)}}{2} \tag{55}$$

which is always positive provided the rates in $P$ are positive, and

$$\lambda_2 = \frac{p_{11} + p_{22} - \sqrt{(p_{11} + p_{22})^2 - 4(p_{11}p_{22} - p_{12}^2)}}{2} \tag{56}$$

which is negative (as needed for stability) if and only if $p_{12}^2 > p_{11}p_{22}$.

Altogether we have the following *coexistence conditions* for a symmetric two-species system with equal local extinction rates:

$$p_{12} > \max(p_{11}, p_{22}) \tag{57}$$

and

$$m < \frac{p_{12}^2 - p_{11}p_{22}}{2p_{12} - p_{11} - p_{22}} \,. \tag{58}$$

The first condition is derived by combining the feasibility requirements $y_1^* > 0$ and $y_2^* > 0$ with the local stability condition $p_{12}^2 > p_{11}p_{22}$. In fact, because any symmetric $2 \times 2$ matrix has constant off-diagonal elements, the analysis in SI Species-Specific Memory Effects proves that this condition (Eq. 57) is sufficient for global stability of the coexistence equilibrium. This is true even when $m_1 \neq m_2$.

The second condition (Eq. 58) arises from $1 > y_1^* + y_2^*$.

The simplicity of the two-species community allows an intuitive interpretation of these results. Examining the formulas for equilibrium frequencies (Eqs. 50 and 53), one can see that changing each parameter value has sensible effects. For example, increasing $m_1$, the local extinction rate of species 1, decreases $y_1^*$ and $x_1^*$ relative to $y_2^*$ and $x_2^*$, respectively. Similarly, increasing $p_{11}$, the rate at which species 1 colonizes patches last occupied by conspecifics, increases both $y_1^*$ and $x_1^*$, while increasing $p_{12}$ increases $x_1^*$ relative to $x_2^*$.

The stability condition for symmetric systems has the natural interpretation that each species must be better at colonizing the other species' patches than its own, generating negative frequency dependence. When this condition is met, it is possible for two species to survive together where either would go extinct individually. This is a simple example of the general feasibility-robustness relationship discussed in Sections 2.C and 3.C. For a single species, our model reduces to the classic Levins metapopulation model, and the lone species persists if $p > m$. Now consider two species, each with (species-specific) colonization rates $p$ in isolation. When these species are present together in a landscape, with $p_{12} = p_{21} > p$, they can withstand local extinction rates up to $\frac{p+p_{12}}{2}$ (following Eq. 58), which is always larger than $p$ – possibly substantially so. For the range $p < m < \frac{p+p_{12}}{2}$, the two-species community will persist, but either species alone would not. A similar (but somewhat more complicated) beneficial effect of diversity is present whenever the matrix $P$ is symmetric, even as $p_{11}$, $p_{22}$, $m_1$, and $m_2$ vary.

Each of these results extends in some way to more diverse communities, but not straightforwardly. For example, with more than two species, the effects of each parameter on the equilibrium frequencies can be complex and indirect. In these cases, it is not possible to solve for the $x^*$ values analytically except in special cases. And while the intuition behind the stability condition found here for symmetric two-species systems extends broadly, the quantitative condition itself does not. The condition $p_{ij} < \max(p_{ii}, p_{jj})$ is neither necessary nor sufficient for coexistence in many-species communities. Instead, it is necessary to consider the eigenvalues of $P$, as captured by Eq. 39.

### 4 Nonsymmetric Memory Effects

When $P$ is no longer symmetric, we are unable to find a general characterization of $\boldsymbol{m}$ and $P$ yielding coexistence. Many behaviors are possible, including non-point attractors. While a complete picture of the model dynamics awaits further study, we consider two special (but nonsymmetric) structures that are potentially biologically relevant and amenable to closer study. In subsection Simulating Dynamics for Random Nonsymmetric Memory Effects, we also present the outcomes of numerical simulations with many nonsymmetric colonization rate matrices sampled at random. These results illustrate general patterns in the dynamics across parameter space.

#### 4.1 Symmetrizable matrices

In SI Symmetric Memory Effects, we implicitly assumed equal dispersal rates in defining the rates $p_{ij}$, which in fact represent a composite of two factors: dispersal (ability to reach a new patch) and establishment (ability to successfully establish residence). If each species has a characteristic dispersal rate, $c_i$, and now we assume the establishment rates $p'_{ij}$ are symmetric, then $p_{ij} = c_i\, p'_{ij}$, and $P$ is called *symmetrizable*. This kind of matrix is nonsymmetric, but can be made symmetric by pre-multiplication with a diagonal matrix. Extensive simulations suggest that stability in this case is controlled entirely by $P'$, the matrix of establishment rates. In particular, to examine the consequences of variation in $\boldsymbol{c}$, we drew random symmetric $P$ matrices with $n = 4$ and $p'_{ij} = p'_{ji} \sim U[0, 1]$ and numerically checked local stability of the coexistence equilibrium for different choices of $\boldsymbol{c}$. This procedure followed closely the analysis of variation in $\boldsymbol{m}$, discussed in SI Symmetric Memory Effects. To assess whether the stability criterion derived with equal dispersal rates remained necessary and sufficient for stability in this generalized setting, we used 1000 randomly distributed $P'$ matrices with exactly one positive eigenvalue (putatively stable) and 1000 matrices with additional positive eigenvalues (putatively unstable). For each matrix, we considered 1000 random $\boldsymbol{c}$ vectors. To avoid sampling many choices of $\boldsymbol{c}$ incompatible with feasibility, which would be computationally costly, we first sampled a vector $\boldsymbol{l}$ uniformly from the simplex, which we took to be proportional to $\boldsymbol{y}^*$. Then we computed $D(\boldsymbol{c})^{-1}$ as $P'\boldsymbol{l}$ and chose $m$ (equal for all species) uniformly distributed between 0 and 1. Finally, the equilibrium frequencies were computed accordingly. This sampling strategy is commonly used to draw random parameters compatible with feasibility (see, for example, [16, 17, 18]). Across all $10^6$ combinations of $P'$ and $\boldsymbol{c}$ for each stability category (stable or unstable), we found that the stability of the system with variation in $c_i$ was always correctly predicted by the stability of the associated system with no variation. This suggests that the condition $P'$ *has exactly one positive eigenvalue* is necessary and sufficient for stability of any community with symmetric memory effects, regardless of the values of each species' colonization ability.

R scripts implementing these simulations are available on GitHub [19].

#### 4.2 Successional cycles

One tractable special case is when $P$ takes the form of a cyclic permutation matrix. As discussed in the main text, this choice of $P$ can be viewed as a simple model for successional dynamics. And we will see that this case illustrates some general features for nonsymmetric $P$, as well.

A cyclic permutation matrix can be written as

$$Q = \begin{pmatrix} 0 & 0 & \dots & 0 & 1 \\ 1 & 0 & \dots & 0 & 0 \\ 0 & \ddots & \ddots & \vdots & \vdots \\ \vdots & \ddots & \ddots & 0 & 0 \\ 0 & \dots & 0 & 1 & 0 \end{pmatrix} \tag{59}$$

assuming species are labeled in the appropriate order. When $n = 3$, this yields the well-known "rock-paper-scissors" dynamics. The key property of this kind of matrix, for our purposes, is that $Q^{-1} = Q^T$. We will use this fact to write the equilibrium frequencies and study the local stability of the coexistence equilibrium.

We assume that all species have identical local extinction rates $m$ and colonization rates $c$ (i.e., $P = cQ$) for simplicity. The equilibrium frequencies for a community of $n$ species are then

$$\begin{aligned} y_i^* &= \frac{m}{c} \\ x_i^* &= \frac{1}{n} - \frac{m}{c}, \quad \text{for all } i. \end{aligned} \tag{60}$$

The only condition for feasibility in this case is that $m < \frac{c}{n}$. Although $P$ is nonsymmetric, the equivalence of species in this highly idealized case means that $\boldsymbol{y}^*$ and $\boldsymbol{x}^*$ are both constant, and therefore proportional. Abusing notation slightly, we let $k = \frac{c}{mn} - 1$ be the constant of proportionality.

The Jacobian evaluated at the coexistence equilibrium is given by

$$J^* = \begin{pmatrix} 0 & k\frac{m}{c}P \\ mI - \frac{m}{c}P^T & -kmI \end{pmatrix} \tag{61}$$

which yields the system of equations

$$\begin{aligned} k\frac{m}{c}P\boldsymbol{v}_i &= \lambda_i \boldsymbol{u}_i \\ m\boldsymbol{u}_i - \frac{m}{c}P^T\boldsymbol{u}_i - km\boldsymbol{v}_i &= \lambda_i \boldsymbol{v}_i \end{aligned} \tag{62}$$

for the $i$th eigenvalue and eigenvector of $J^*$. Using the fact that $P^T = c^2P^{-1}$, and assuming $\lambda_i \neq 0$, we can re-arrange Eq. 62 to obtain

$$P\boldsymbol{v}_i = (c + \frac{c}{m}\lambda_i + \frac{c}{km^2}\lambda_i^2)\boldsymbol{v}_i \tag{63}$$

which implies that the eigenvalues of $J^*$ and $Q$ are related by

$$\lambda(Q)_i = 1 + \frac{1}{m}\lambda_i + \frac{1}{km^2}\lambda_i^2. \tag{64}$$

Solving for $\lambda_i$, we find

$$\lambda_i = \frac{-km\left(1 \pm \sqrt{1 - \frac{4}{k}(1 - \lambda(Q)_i)}\right)}{2}. \tag{65}$$

The quantity $km$ is always positive when the coexistence equilibrium is feasible, so the real parts of the $\lambda_i$ will be all negative if and only if $1 > \text{Re}\left(\sqrt{1 - \frac{4}{k}(1 - \lambda(Q)_i)}\right)$ for every $\lambda(Q)_i$. In general, $\lambda(Q)_i$ may be complex, so we write these eigenvalues more explicitly as $\lambda(Q)_i = a + bi$. The real part of a complex square root then gives us

$$\begin{aligned} 1 > \text{Re}\left(\sqrt{1 - \frac{4}{k}(1 - \lambda(Q)_i)}\right) \\ = \sqrt{\frac{1 - \frac{4}{k} + \frac{4}{k}a + \sqrt{(1 - \frac{4}{k} + \frac{4}{k}a)^2 + (\frac{4}{k}b)^2}}{2}} \\ = \sqrt{\frac{1 - \frac{4}{k} + \frac{4}{k}a + \sqrt{1 - \frac{8}{k}(1 - \frac{4}{k})(1 - a)}}{2}} \end{aligned} \tag{66}$$

using the fact that $a^2 + b^2 = 1$ for any permutation matrix to obtain the last line. This expression can be simplified substantially to produce the equivalent inequality

$$k > a + 1 \tag{67}$$

or

$$m < \frac{c}{n(a+2)}. \tag{68}$$

Finally, we consider the eigenvalues of $Q$, which are the $n$th roots of unity, with real parts $a_k = \cos\left(\frac{2\pi k}{n}\right)$ for $k = 0, ..., n-1$. For $k = 0$, the associated $\lambda_i$ is the (structural) zero eigenvalue of $J$. The relevant bound for $m$ is found by considering the largest (non-trivial) value of $a_k$, which is $\cos\left(\frac{2\pi}{n}\right)$, for $k = 1$. This gives us the stability threshold

$$m_c = \frac{c}{n\left(\cos\left(\frac{2\pi}{n}\right) + 2\right)}. \tag{69}$$

When $m < m_c$, the coexistence equilibrium is stable. For $n > 2$, this threshold is always less than the threshold for feasibility, $m_{\max} = \frac{c}{n}$. For the range of $m$ values in between, $m_c < m < m_{\max}$, we observe limit cycles (Fig. S2). Numerical evidence shows that the amplitude of these cycles grows rapidly as $m$ increases, so that for $n \geq 3$, all species go extinct even before $m_{\max}$ is reached (Fig. S3). The parameter space where stable coexistence is possible shrinks rapidly as $n$ increases; for large $n$, we have $m_c \approx \frac{c}{3n}$. As one might expect, higher colonization rates increase the stability threshold, as well as the feasibility threshold. Both are proportional to $c$, so that the stable fraction of feasible parameter space has no dependence on $c$.

##### 4.3 Simulating dynamics for random nonsymmetric memory effects

The previous sections suggest that even keeping $m$ equal for all species, communities with nonsymmetric memory effects may behave similarly to the symmetric case, or very dissimilarly, with the possibility for limit cycles and stability dependent on the magnitude of $m$. To better understand which outcomes might be typical, we sampled nonsymmetric $P$ matrices and integrated the model dynamics for a range of $m$ values (equal for all species) up to $m_{\max}$

for each model community. Specifically, we sampled $P$ matrices for 3-species communities with $p_{ij} \sim U[0,1]$ i.i.d. and discarded matrices which were not potentially feasible (i.e., $P^{-1}\mathbf{1}$ not all positive). To save computing time, we checked whether each matrix had a locally stable equilibrium for $m$ small ($m = \frac{1}{100}m_{\max}$); if not, it was discarded. This procedure was motivated by the hypothesis that if a $P$ matrix is compatible with stability for some choice of $m$, say $m'$, then we will find stability for any $m < m'$. This hypothesis is suggested by the case of cyclic $P$, and supported by the simulation results. However, to ensure the observed pattern was a not a consequence of selecting communities stable at low $m$, we also repeated the simulations with $P$ drawn as before, but checking local stability for $m$ large ($m = \frac{99}{100}m_{\max}$). For both sets of simulations, we collected 200 matrices meeting the stated criteria, and then simulated the model dynamics for 50 values of $m$ evenly spaced between 0 and $m_{\max}$. Each community was initialized at a random point near the coexistence equilibrium, to avoid long transients. We integrated the dynamics for 5000 time steps, and then checked if (i) species frequencies had converged to the coexistence equilibrium, (ii) one or more species had gone extinct, or (iii) the dynamics had converged to a stable limit cycle (using the criterion that the final frequencies had been visited at least 5 times independently through the dynamics). If none of these criteria were met, we integrated another 5000 time steps and checked again. This was repeated up to 10 times, until an outcome could be classified.

The results of these simulations are shown in Fig. S4. We find that a progression from stability, to limit cycles, to instability as $m$ increases is a common outcome (47% of the communities lose stability). Thus, the qualitative behavior shown analytically for cyclic $P$ seems to be a fairly general feature of the dynamics when $P$ is nonsymmetric. However, many communities exhibited a stable equilibrium for all values of $m$, consistent with the symmetric and symmetrizable cases. When we required $P$ to produce a stable equilibrium at high $m$ (Figure S4, right), the only observed outcome was stability for all values of $m$. Together, these results strongly support our hypothesis that stability can only be lost, not gained, as $m$ increases.

### 5 Waning Memory Effects

#### 5.1 Modified model

To relax the assumption that patches remain in state $i$ indefinitely after occupation by species $i$, we extended our model to include an additional "naïve" state. The dynamics of this extended model are given by:

$$\begin{aligned}
\frac{dx_i(t)}{dt} &= -m_i x_i(t) + c_i x_i(t) z(t) + x_i(t) \sum_{j=1}^{n} p_{ij} y_j(t) \\
\frac{dy_i(t)}{dt} &= m_i x_i(t) - d_i y_i(t) - y_i(t) \sum_{j=1}^{n} p_{ji} x_j(t) \\
\frac{dz(t)}{dt} &= \sum_{j=1}^{n} d_j y_j(t) - z(t) \sum_{j=1}^{n} c_j x_j(t)
\end{aligned} \tag{70}$$

where $z(t)$ is the frequency of naïve patches. All parameters are interpreted as before, with the addition of decay rates $d_i$ (the rate at which patches in state $i$ transition into the naïve state) and parameters $c_i$, which encode the rate at which species $i$ colonizes naïve patches. While this model easily accommodates differences in the durability of memory effects or baseline colonization rates across species, we will assume all species have equal demographic rates $m$, $c$, and $d$ for simplicity. We also focus on the case where $P$ is symmetric. In this section, we show that the stability condition derived for symmetric $P$ without memory extends straightforwardly to the modified model.

For the coexistence equilibrium of the modified model one finds

$$\begin{aligned} \boldsymbol{y}^* &= (m - \frac{d}{k})\, P^{-1}\mathbf{1} \\ \boldsymbol{x}^* &= k\,\boldsymbol{y}^* \\ z^* &= \frac{d}{c\,k}. \end{aligned} \tag{71}$$

Just as before, $\boldsymbol{x}^*$ is proportional to $\boldsymbol{y}^*$, with constant of proportionality $k$. Now, however, we have the modified zero-sum condition $\mathbf{1}^T(\boldsymbol{x}^* + \boldsymbol{y}^* + z^*) = 1$, so $k$ satisfies the quadratic equation

$$(m - \frac{d}{k})\frac{1}{q} + (k\,m - d)\frac{1}{q} + \frac{d}{c\,k} = 1 \tag{72}$$

or

$$k^2 + (1 - \frac{d}{m} - \frac{q}{m})\,k + \frac{d}{m}(\frac{q}{c} - 1) = 0 \tag{73}$$

where $q = (\mathbf{1}^T P^{-1}\mathbf{1})^{-1}$. The value $q$ can be seen as a statistic summarizing the effective whole-community colonization rate, in the following sense: If matrix $P$ has summary statistic $q$, then the associated system behaves identically – with respect to its "demographic" features, such as $k$ values and demographic eigenvalues (see below) – to a single-species metapopulation with colonization rate $q$.

From Eq. 73, there are two solutions for $k$ (and possibly two distinct equilibria), given by

$$k = \frac{\frac{d}{m} + \frac{q}{m} - 1 \pm \sqrt{(1 - \frac{d}{m} - \frac{q}{m})^2 - 4\frac{d}{m}(\frac{q}{c} - 1)}}{2}. \tag{74}$$

An examination of Eq. 71 shows that for feasibility, we must have all entries of $P^{-1}\mathbf{1} > 0$, as before, and also $k > \frac{d}{m}$. The latter inequality might be satisfied by 0, 1 or 2 values of $k$. When $c > m$, there will be exactly one such $k$. On the other hand, when $d + m < q$, there will be no $k$ compatible with feasibility. These conditions are found by substituting $k = k' + \frac{d}{m}$ into Eq. 73 and applying Descartes' rule of signs. In cases where $c < m$ and $d + m > q$, there may be 0 or 2 feasible values of $k$, depending on the sign of the discriminant in Eq. 74. The condition $c < m$ implies that the zero-biomass equilibrium, $z^* = 1$, is attractive, and we will see that the second equilibrium, corresponding to the smaller value of $k$, is always unstable when feasible. These properties signal the presence of a strong Allee effect. If the coexistence equilibrium corresponding to the larger value of $k$ is locally stable in this case, there will be bistability, with trajectories attracted to the zero-biomass equilibrium or the coexistence equilibrium depending on the initial condition.

When is the coexistence equilibrium stable in this extended model? Evaluating the Jacobian at the equilibrium corresponding to either $k$ (and assuming feasibility), we find

$$J^* = \begin{pmatrix} 0 & D(\boldsymbol{x}^*)P & c\,\boldsymbol{x}^* \\ D(\boldsymbol{m}) - D(\boldsymbol{y}^*)P & -D(P\boldsymbol{x}^*) - d\,I & 0 \\ -c\,z^*\,\mathbf{1}^T & d\,\mathbf{1}^T & -c\,(\mathbf{1}^T\boldsymbol{x}^*) \end{pmatrix} \tag{75}$$

with appropriate $\boldsymbol{x}^*, \boldsymbol{y}^*$, and $z^*$.

As in SI Symmetric Memory Effects, we can use an equilibrium relationship $D(\boldsymbol{m})\boldsymbol{x}^* = D(\boldsymbol{y}^*)(P\boldsymbol{x}^* - dI)$ to re-write this matrix slightly:

$$J^* = \begin{pmatrix} 0 & D(\boldsymbol{x}^*)P & c\,\boldsymbol{x}^* \\ D(\boldsymbol{m}) - D(\boldsymbol{y}^*)P & -kmI & 0 \\ -c\,z^*\,\mathbf{1}^T & d\,\mathbf{1}^T & -c\,(\mathbf{1}^T\boldsymbol{x}^*) \end{pmatrix}. \tag{76}$$

Let us denote the upper-left $2n \times 2n$ block of $J^*$ by $Q$. Notice that $Q$ is of exactly the same form as Eq. 32. We will see that $2n - 2$ of the eigenvalues of $J$ are also eigenvalues of $Q$, governed by the same stability criterion as in the case without waning memory. In other words, the condition *$P$ has exactly one positive eigenvalue* is now a necessary condition for local stability of the coexistence equilibrium. In fact, we will see that it is also sufficient.

But first we will consider the eigenvalues of $J^*$ corresponding to those of $Q$ more carefully. Any eigenvectors of $Q$, written with components of length $n$ as $(\boldsymbol{u}, \boldsymbol{v})^T$, must satisfy the system

$$\begin{aligned} D(\boldsymbol{x}^*)P\boldsymbol{v} &= \lambda\boldsymbol{u} \\ m\boldsymbol{u} - D(\boldsymbol{y}^*)P\boldsymbol{u} - km\boldsymbol{v} &= \lambda\boldsymbol{v}. \end{aligned} \tag{77}$$

Multiplying both equations by $\mathbf{1}^T$, we obtain

$$\begin{aligned} (km - d)(\mathbf{1}^T\boldsymbol{v}) &= \lambda(\mathbf{1}^T\boldsymbol{u}) \\ m(\mathbf{1}^T\boldsymbol{u}) - (m - \frac{d}{k})(\mathbf{1}^T\boldsymbol{u}) - km(\mathbf{1}^T\boldsymbol{v}) &= \lambda(\mathbf{1}^T\boldsymbol{v}) \end{aligned} \tag{78}$$

and so

$$km(\mathbf{1}^T\boldsymbol{v}) = \frac{d}{k}(\mathbf{1}^T\boldsymbol{u}) - \lambda(\mathbf{1}^T\boldsymbol{v}) = \lambda(\mathbf{1}^T\boldsymbol{u}) + d(\mathbf{1}^T\boldsymbol{v}). \tag{79}$$

This last set of equalities can be satisfied by $(\mathbf{1}^T\boldsymbol{u}) = (\mathbf{1}^T\boldsymbol{v}) = 0$, or when $(\mathbf{1}^T\boldsymbol{u}), (\mathbf{1}^T\boldsymbol{v}) \neq 0$. In the latter case, one can divide by $(\mathbf{1}^T\boldsymbol{u})$ to obtain the system

$$\begin{aligned} (d - km)w^2 - kmw + \frac{d}{k} &= 0 \\ \lambda &= \frac{d}{k}\frac{1}{w} - km \end{aligned} \tag{80}$$

for $w = (\mathbf{1}^T \boldsymbol{v})/(\mathbf{1}^T \boldsymbol{u})$ and the eigenvalue, $\lambda$. Two eigenvalues of $Q$ are associated with Eq. 80, and the remaining $2n-2$ must be associated with eigenvectors that satisfy $(\mathbf{1}^T \boldsymbol{u}) = (\mathbf{1}^T \boldsymbol{v}) = 0$. The latter eigenpairs of $Q$ generate eigenpairs of $J^*$ with the same eigenvalues and augmented eigenvectors $(\boldsymbol{u}\,, \boldsymbol{v},\, 0)^T$. This is easy to verify:

$$J^*\,(\boldsymbol{u}\,,\boldsymbol{v}\,,0)^T = \begin{pmatrix} Q & \cdots & c\,\boldsymbol{x}^* \\ \vdots & \ddots & 0 \\ -c\,\boldsymbol{z}^*\,\mathbf{1}^T & d\,\mathbf{1}^T & -c\,(\mathbf{1}^T\boldsymbol{x}^*) \end{pmatrix} (\boldsymbol{u}\,,\boldsymbol{v}\,,0)^T$$
$$= \begin{pmatrix} \lambda\,(\boldsymbol{u}\,,\boldsymbol{v})^T + 0 \\ -c\,\boldsymbol{z}^*\,\mathbf{1}^T\boldsymbol{u} + d\,\mathbf{1}^T\boldsymbol{u} + 0 \end{pmatrix} = \begin{pmatrix} \lambda\,(\boldsymbol{u}\,,\boldsymbol{v})^T \\ 0 \end{pmatrix} = \lambda\,(\boldsymbol{u}\,,\boldsymbol{v}\,0)^T. \tag{81}$$

Since $Q$ is of the same form as Eq. 32, its eigenvalues (all $2n$) are related to the eigenvalues of $S = D(\boldsymbol{y}^*)^{1/2} P D(\boldsymbol{y}^*)^{1/2}$ according to Eq. 39. The Perron eigenvector of $S$ is non-negative, and therefore must violate $(\mathbf{1}^T \boldsymbol{u}) = (\mathbf{1}^T \boldsymbol{v}) = 0$, so the two eigenvalues of $Q$ that are not also eigenvalues of $J^*$ are those associated with the Perron eigenvalue of $S$. This eigenvalue is equal to $m - \frac{d}{k}$, which bounds the spectral radius of $S$ below $m$, and following the logic developed in the unmodified case, the remaining $2n-2$ eigenvalues of $Q$ will be negative if and only if $P$ has exactly exactly one positive eigenvalue.

Finally we consider the remaining three eigenvalues of $J^*$. One of these must be zero, because the dynamics are confined to the simplex. We call the final two eigenvalues of $J^*$ its "demographic" eigenvalues, as they depend only on $c$, $m$, $q$, and $d$ – and not on the structure of $P$. Each of these three eigenvalues correspond to eigenvectors of the form $(\boldsymbol{x}^*, a_1\boldsymbol{x}^*, a_2)^T$ for undetermined constants $a_1$ and $a_2$. Assuming this structure and multiplying by $J^*$ yields the nonlinear scalar system

$$\begin{aligned} a_1\,k\,(m - \frac{d}{k}) + a_2\,c &= \lambda \\ \frac{d}{k} - a_1\,k\,m &= a_1\,\lambda \\ (\frac{-d}{k} + a_1\,d - a_2\,c)(k\,m - d)\frac{1}{q} &= a_2\,\lambda. \end{aligned} \tag{82}$$

We solved this system using Wolfram Mathematica (12.2) (scripts available on GitHub [19]). One solution is $\lambda = 0$, as expected. The two demographic eigenvalues are given by

$$\frac{rd - km(r+1) \pm \sqrt{(rd - km(r-1))^2 + 4d(\frac{d}{k} - m)(r-1)}}{2}. \tag{83}$$

using the shorthand $r = \frac{c}{q}$. We used Mathematica to verify that these eigenvalues are always negative for the unique feasible coexistence equilibrium (when $c > m$). When two equilibria are feasible, we verified that one or both eigenvalues are always positive for the equilibrium corresponding to smaller $k$. Although we are unable to prove it analytically, extensive numerical simulations suggest that the other equilibrium is always stable in this case, as well.

Altogether, this analysis suggests that if the equilibrium corresponding to larger $k$ is feasible, it will always be locally stable provided $P$ has exactly one positive eigenvalue. If the second equilibrium is also feasible, the model exhibits bistability. The possible outcomes across parameter space are illustrated graphically in Fig. 5 in the main text.

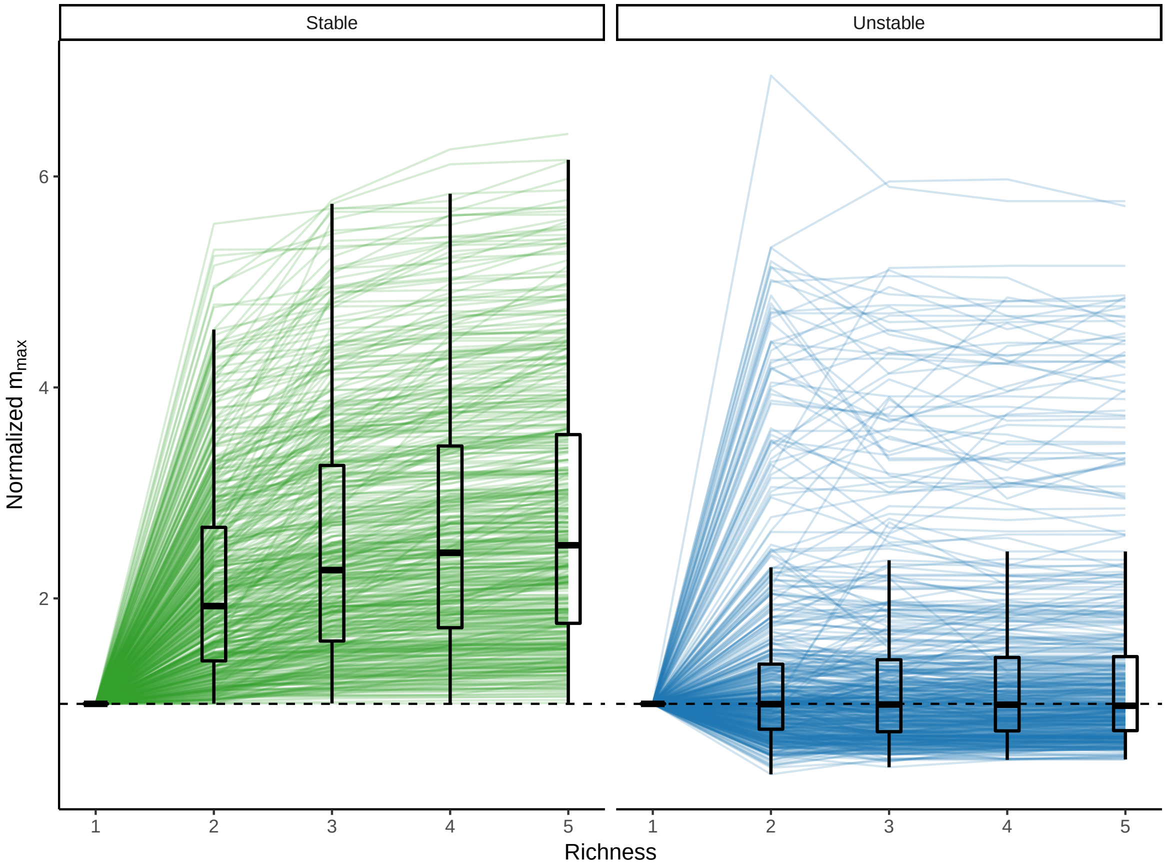


Figure 1: Coexistence criteria induce a positive relationship between diversity and robustness. Along 500 random assembly sequences leading to coexisting communities of 5 species, $m_{\max}$ always increases (left). When the final community does not coexist, $m_{\max}$ is never strictly increasing. Here we show a normalized measure of $m_{\max}$ for each assembly sequence, obtained by dividing $m_{\max}$ at each level of richness by its value for the first species alone. See SI Symmetric Memory Effects for simulation details.

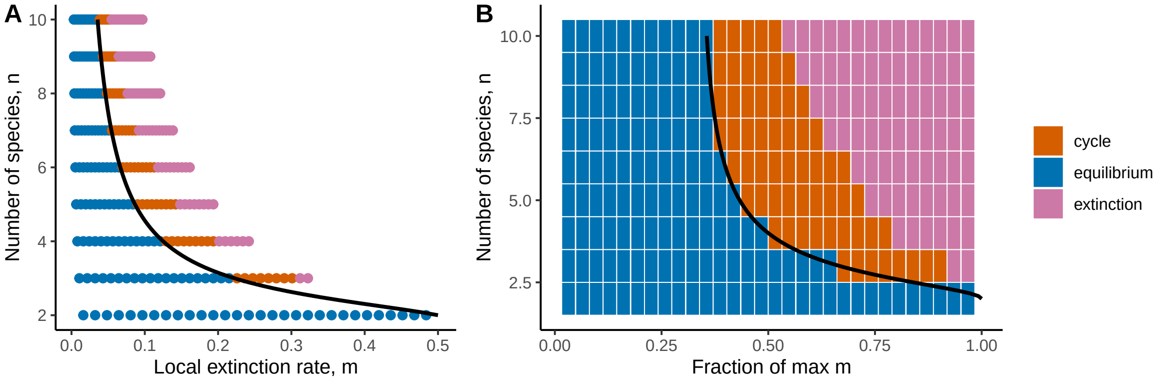


Figure 2: Long-term dynamics for cyclic memory effects. When $P$ is proportional to a cyclic permutation matrix, all species may coexist stably, cycle, or go extinct, depending on the magnitude of $m$. (A) Each point represents the outcome of simulated dynamics with $n$ species and local extinction rate $m$. Coexistence (blue) occurs for sufficiently small values of $m$. Above a threshold given by Eq. 69 (black curve), stability is lost, leading to limit cycles (orange) and eventually extinction of all species (pink). For each $n$, we simulated communities at 30 values of $m$ evenly space between 0 and $m_{\max}$. Beyond $m_{\max}$, there is no feasible equilibrium. Notice that the feasibility threshold declines sharply with $n$. (B) As in (A), except the x-axis shows $m$ values normalized by $m_{\max}$, for each $n$ (rows).

[3] Wolfgang Hahn, Hans H Hosenthien, and H Lehnigk. *Theory and application of Liapunov's direct method*, volume 3. Prentice-Hall Englewood Cliffs, NJ, 1963.

[4] Robert MacArthur. Species packing and competitive equilibrium for many species. *Theoretical population biology*, 1(1):1–11, 1970.

[5] Bo S Goh. Global stability in many-species systems. *The American Naturalist*, 111(977):135–143, 1977.

[6] Léon Brenig. Complete factorisation and analytic solutions of generalized lotka-volterra equations. *Physics Letters A*, 133(7-8):378–382, 1988.

[7] Benito Hernández-Bermejo, Víctor Fairén, and Léon Brenig. Algebraic recasting of nonlinear systems of odes into universal formats. *Journal of Physics A: Mathematical and General*, 31(10):2415, 1998.

[8] Gabor Szederkenyi, Attila Magyar, and Katalin M Hangos. *Analysis and control of polynomial dynamic models with biological applications*. Academic Press, 2018.

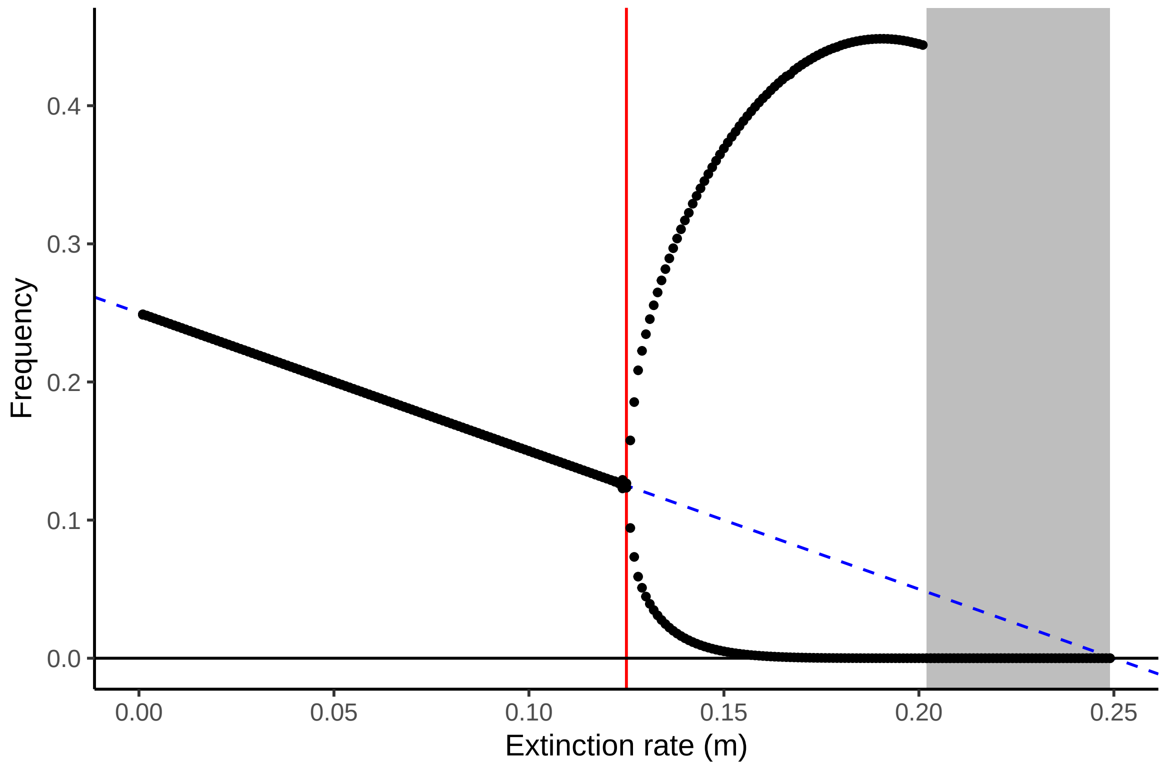


Figure 3: Bifurcation diagram for cyclic $P$. As $m$ increases, the (common) equilibrium abundance of all species decreases linearly. At a threshold indicated in red ($m_c$), there is a bifurcation point where the dynamics begin to cycle (minimum and maximum points shown). The amplitude of these cycles increases rapidly, leading to the extinction of all species before the feasibility threshold is reached at $m_{\max} = 0.25$ (gray shaded area). In a community experiencing gradually increasing $m$ (e.g. increasing disturbance), predicting future equilibrium frequencies based on the rate of decline at small $m$ (dashed blue line) fails suddenly at $m = m_c$. These simulations used $n = 4$ and $c = 1$, but qualitatively identical behavior is observed for any $n \geq 3$ and any $c$.

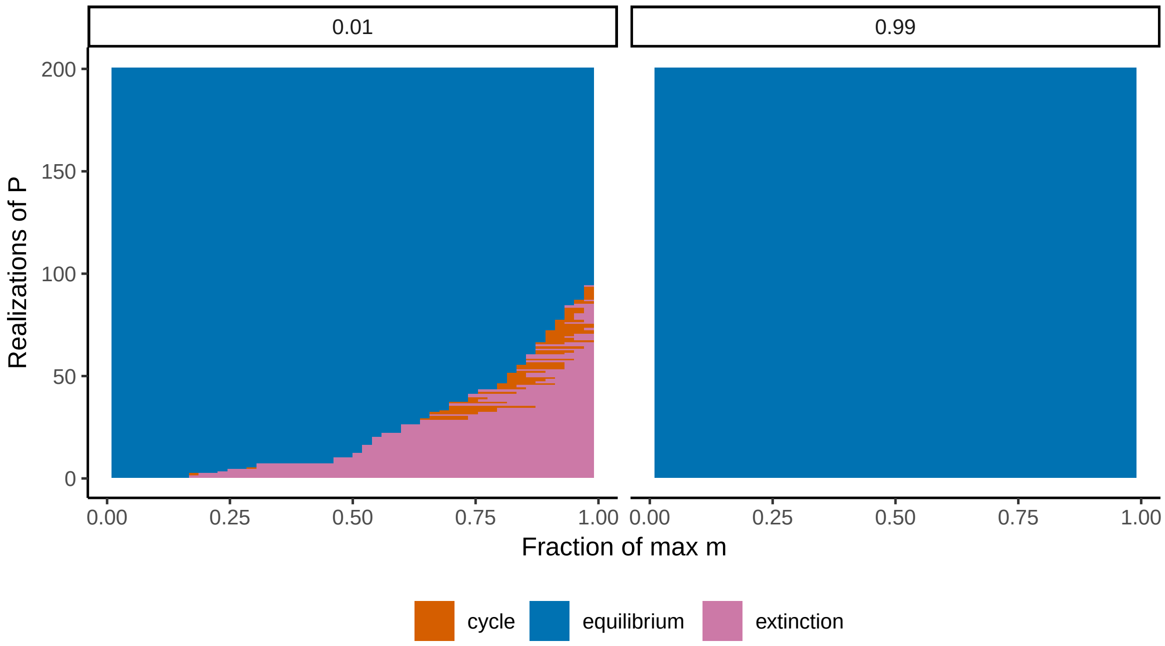


Figure 4: Long-term dynamics for nonsymmetric $P$. Each row shows the long-term dynamics for a different 3-species community with randomly-distributed $P$, along a gradient of $m$ values. Communities were selected to be stable at either low ($m = \frac{1}{100} m_{\max}$; left) or high ($m = \frac{99}{100} m_{\max}$; right) values of $m$. For the latter, stability may be lost as $m$ increases (this occurs in 47% of cases shown). As suggested by the analysis of cyclic $P$, there is often a narrow region where limit cycles are observed between loss of stability of the equilibrium and the extinction of some species. When $P$ is selected for stability at high $m$ (right), stability is never lost. In the left panel, communities are ordered along the y-axis by the point at which they lose stability.
